# Measuring cell-to-cell expression variability in single-cell RNA-sequencing data: a comparative analysis and applications to B cell ageing

**DOI:** 10.1101/2022.11.24.517880

**Authors:** Huiwen Zheng, Jan Vijg, Atefeh Taherian Fard, Jessica Cara Mar

## Abstract

**Background:** Single-cell RNA-sequencing (scRNA-seq) technologies enable the capture of gene expression heterogeneity and consequently cell-to-cell variability at the cell type level. Although different methods have been proposed to quantify cell-to-cell variability, it is unclear what the optimal statistical approach is, especially in light of challenging data structures that are unique to scRNA-seq data like zero inflation.

**Results:** In this study, we conducted a systematic evaluation of cell-to-cell gene expression variability using 14 different variability metrics that are commonly applied to transcriptomic data. Performance was evaluated with respect to data-specific features like sparsity and sequencing platform, biological properties like gene length, and the ability to recapitulate true levels of variability based on simulation and known biological gene sets like ribosomal genes and stably expressed genes. scran had the strongest all-round performance, and this metric was then applied to investigate the changes in cell-to-cell variability that occur during ageing. Studying ageing showcases the value of cell-to-cell variability as it is a genetically-regulated program that is influenced by stochastic processes.scRNA-seq datasets from hematopoietic stem cells (HSCs) and B lymphocytes and other cell types from this differentiation lineage were used with scran to identify the genes with consistent patterns of variable and stable expression profiles during differentiation. Furthermore, to understand the regulatory relationship for genes that were differentially-variable in their expression between young and old mice, we constructed networks using transcription factors and their known targets for HSC and B lymphocyte cells. Comparisons of these networks identified a shared TF *Sfpi1* that although was seen to increase in gene expression variability in old mice versus young in both cell types, the corresponding targets were distinct and their gene expression variability had different directions between cell types.

**Conclusions:** Through these analyses, we highlight the importance of capturing cell-to-cell gene expression variability in a complex biological process like differentiation and ageing, and emphasise the value and specificity of interpreting these findings at the level of individual cell types.

## Background

Cells are the basic units of life, and heterogeneity in gene expression is something that exists between cells, even in populations of genetically identical cells. At the molecular level, cell-to-cell variability results from cells that cancel out intrinsic noise while amplifying regulated variability [1]. The demonstrated link between cell-to-cell variability and the propagation of gene expression through regulatory networks makes it critically important to model changes in cell-to-cell variability as a means to identify regulators that may have been overlooked by focusing on changes in average expression only.

Advances in single-cell RNA sequencing (scRNA-seq) technologies have pushed the boundaries of the resolution at which gene expression measurements can now be obtained from single cells [2]. The popularity of scRNA-seq datasets has renewed interest in gene expression variability and what this metric can reveal about underlying regulatory processes [3, 4]. Although cell-to-cell variability is a straightforward concept, a range of terms has been used to describe it, including cellular heterogeneity, transcriptional variability, gene expression variability or transcriptional noise. There has been an even greater number of metrics used to measure cell-to-cell variability that follow their own distinct statistical approaches. This is seemingly problematic because each study of cell-to-cell variability adopts its own specific quantitative approach for the analysis. Without an understanding of which variability metrics have the strongest performance for scRNA-seq data it is possible that these studies are using variability estimates in a sub-optimal way, therefore making it difficult to model cell-to-cell variability in complex biological processes such as ageing.

Measuring gene expression variability is challenging for scRNA-seq data because of the typical characteristics that this sequencing approach generates that create additional limitations for modelling transcript counts and the true levels of cell-to-cell variability. For example, sparsity, which is in part driven by stochastic gene expression, makes it challenging to estimate cell-to-cell variability because many traditional summary statistics cannot handle the increased frequency of zeros. Additionally, low mRNA capture efficiency and low sequencing depth contribute to sparsity that can make it challenging to capture the sufficient and equal cell type sizes that are required to model true gene expression variability. On the other hand, genes with very low read counts tend to have more variable expression, resulting in a strong mean-variance relationship [5]. Therefore, if this relationship is not being accounted for, changes in the measured variability may be driven by fluctuations of lowly expressed genes rather than the true variability. While we understand these issues in the context of how we model scRNA-seq data through typical workflows, modelling variability is in some ways more complicated because the variability is more influenced by aspects like sample size, than statistics based on average expression [6].

Studying ageing showcases the value of what can be learned from gene expression variability of scRNA-seq data because ageing is a genetically-regulated program that is executed by a series of cellular and molecular factors and influenced by stochastic processes. For example, bone marrow has been intensively investigated as it is the primary site to produce hematopoietic stem cells (HSCs) which give rise to all the cells involved in the immune response. Throughout life, the function of the bone marrow adapts to match the fluctuating demands of the organism and impacts HSCs self-renewing [7]. The long-term reconstituting HSCs can lead to phenotypically and transcriptionally distinct short-term HSCs that lack durable self-renewal, which can progressively generate lineage-restricted progenitors and mature cells of the myeloid, lymphoid and megaerythroid lineages [8]. Furthermore, the reduction of regenerative capacity of the stem cells and B lymphoid lineages generation rates during ageing result in narrowed clonotypic diversity [9] which presumably must impact cell-to-cell variability of gene expression. These observations highlight the importance of profiling cell-to-cell variability to characterise the developmental process of ageing HSCs and B lymphoid lineages.

Age-dependent changes in gene expression variability are difficult to address because of inherent noise and experimental factors that may influence variability [10]. Large databases like Tabula-Muris-Senis (TMS) [11] and GTEx [12] provide the opportunity to expand our knowledge on how cell-to-cell variability changes during ageing in various organs and tissues. Several studies have reported that gene expression variability increases during ageing such as mouse heart cardiomyocytes [13] and human pancreatic cells [14]. However, these increases in variability during ageing are not consistently observed across all tissue or cell types, e.g. in ageing hematopoietic cell types and mouse brains [15, 16]. These inconsistencies may be due to genuine biological effects, differences in the experimental factors inherent to the design, or the choice of the analysis approach of these datasets. Resolving these inconsistencies to identify what patterns of gene expression variability can be generalised between conditions to help provide insight into complex processes like ageing.

In this study, we conduct a systematic evaluation of a wide range of metrics that are used for measuring gene expression variability in transcriptomic data. The focus of the evaluation was to identify which variability metric had the strongest performance when evaluated against a set of different biological and data-specific features that are relevant to scRNA-seq data. The winner of this benchmarking exercise, scran, was used to investigate the role of cell-to-cell variability in identifying regulatory relationships underlying the ageing process on HSCs and B lymphoid lineages sourced from the TMS database. We observed cellular heterogeneity changes that were cell-type-specific for B lymphoid differentiation during ageing. These results aligned with existing underlying biology and provided additional evidence for stem cell exhaustion and a decline in B lymphomagenesis in ageing. Furthermore, we also developed a new method to identify differentially variable genes in aged HSCs and B cells, which showed distinct regulatory networks with a shared transcription factor. Taken together, our results provide ample evidence that studying cell-to-cell variability at the cell type level has important implications for understanding ageing cells that undergo cell type-specific differentiation processes.

## Results

### Estimating biological variability from scRNA-seq data is influenced by a dataset’s structure

Statistics often perform in a data-dependent manner and the ability to measure biological variability from scRNA-seq data is no exception. Different statistical metrics have been proposed to estimate biological variability and each metric comes with its own strengths and weaknesses. Since these metrics first appeared, the generation of scRNA-seq data and the associated analytical approaches have matured. Collectively, we understand that while datasets can vary widely, there are some structures that scRNA-seq datasets share. This section of our study investigates how specific kinds of biological and data-specific features impact different metrics for estimating biological variability and aims to identify which metric has the best overall performance.

### Investigating how sequencing methods impact metric performance

We assume that a reliable metric for measuring biological variability will estimate this parameter regardless of the sequencing platform used to generate the data. The design of the TMS dataset provided an opportunity to assess how metrics performed under two different sequencing platforms because the same cell types were sequenced with a FACS-sorted Smartseq2 and a 10X Genomics Droplet-based method. These two sequencing methods distinctly capture full-length or 3’ reads, resulting in unique data structures. Our results demonstrated that in general, the platform-specific differences in gene expression variability tend to be larger than the differences due to cell type. These comparisons indicate that platform effects are important to consider when measuring gene expression variability (Figure 1a).

**Figure 1:**
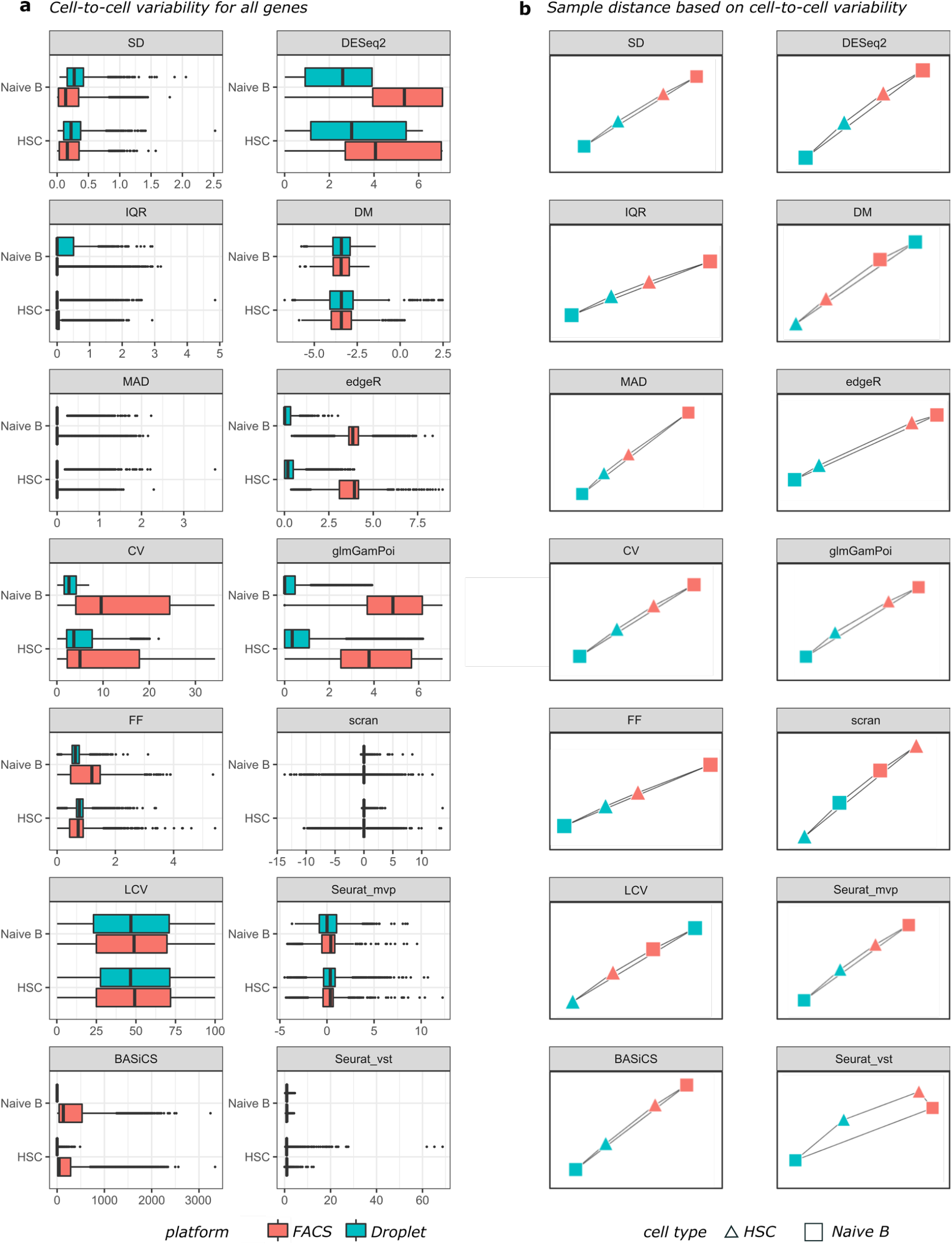
Cell-to-cell variability for all genes measured from 14 metrics between two cell types with two sequencing platforms. a) boxplots for the cell-to-cell variability in each unique sample, coloured by the sequencing platform. b) sample distance between specific cell types under two sequencing platforms calculated from Kolmogorov–Smirnov’s D statistic.

To quantitatively investigate the platform-specific effect, the Kolmogorov–Smirnov’s D statistic was used to measure the distance between the variability metric distributions between the two sequencing methods for the same cell type. We compared these values to the distance between different cells sequenced by the same sequencing method to evaluate the size of the platform-specific effect (Supplementary Table 2). The sample similarities were visualised by the network and the edge lengths were calculated based on the D statistic values (Figure 1b). In general, CV, DESeq2, edgeR and glmGamPoi were impacted most significantly by sequencing methods for both cell types while DM, LCV, scran and Seurat metrics were more robust and showed similar estimated variability within the same cell types regardless of the sequencing method.

It is important to recognise that all sequencing methods have their own degree of technical variability and in fact, each data set comes with its own distribution of expression variability as estimated by each metric. This can be observed through the fact that the distance between the two sequencing methods for the same cell type was not zero for any of the comparisons made in this study (Supplementary Table 2). Therefore, the degree of overall variability even for the same cell type should be handled carefully when analysing data from different sequencing methods.

### Investigating the impact of sample size on metric performance

Increasing the sample size tends to reduce the variance in a population and we investigated metric performance for scRNA-seq data with respect to this criterion. We used the number of cells available for different cell types under the same sequencing method to evaluate the impact of sample size on measuring cell-to-cell gene expression variability, e.g. HSC and naïve B (NB) cells each had a relatively large sample size (∼1000 cells) versus immature B (IB) cells, which had only 50 cells in the TMS dataset (where the cell type sample sizes range from 50 to 1000 cells). Data with small sample sizes (∼50 cells) showed a smaller range in their cell-to-cell variability distribution, especially when using metrics like CV, DESeq2 and edgeR (Figure 2a). Therefore, an insufficient number of cells may lead to biases when calculating the gene expression variability regardless of the metric chosen, resulting in underestimating the amount of cellular heterogeneity, especially in rare cell types.

**Figure 2:**
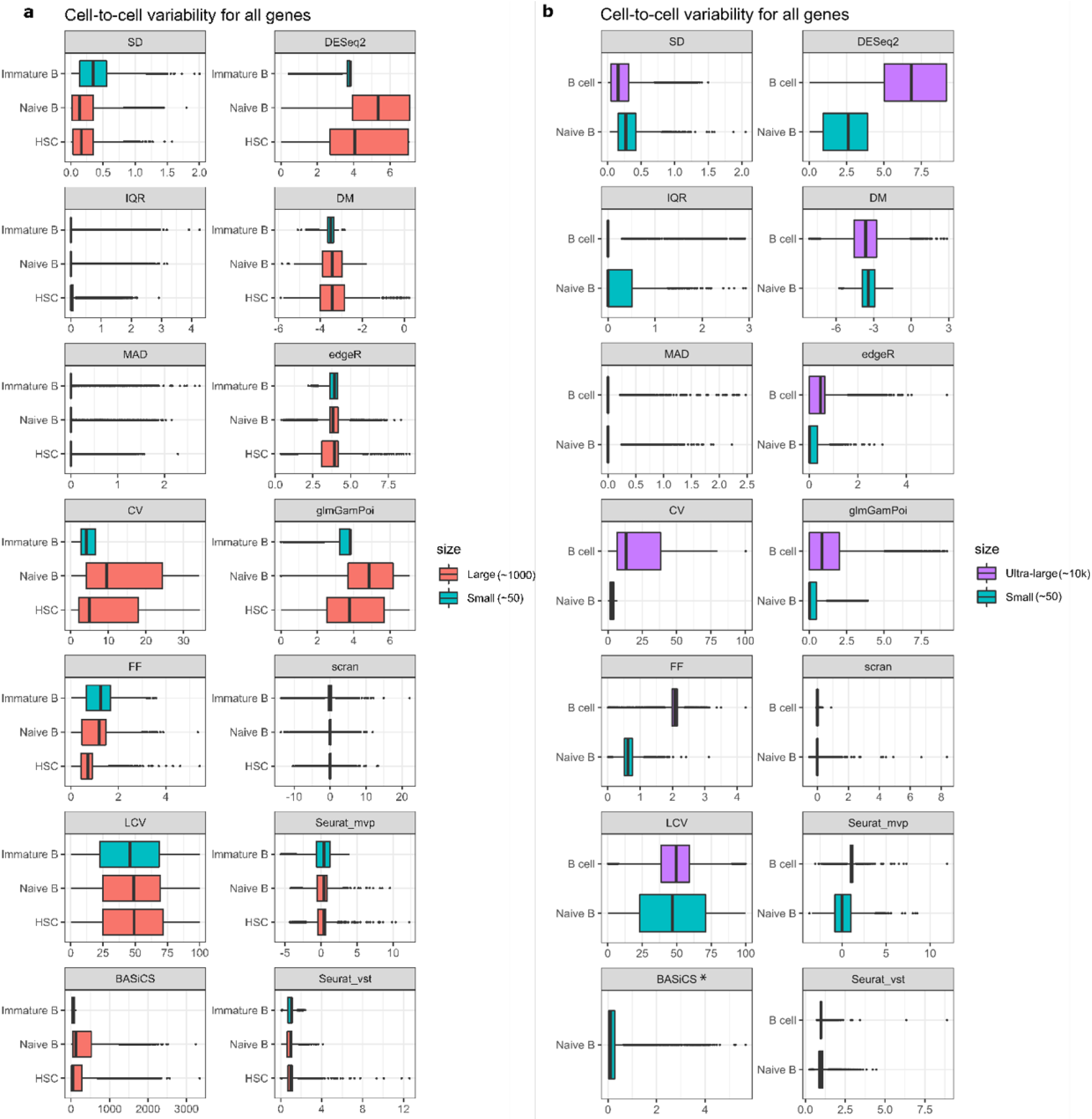
Cell-to-cell variability for all genes measured from 14 metrics among several cell types with different sample sizes on the same sequencing platform. a) boxplots for the cell-to-cell variability in each unique sample, coloured by sample sizes. b) boxplots for cell-to-cell variability for B cells from TMS and an external PBMC sample. *BASiCS was not applied to ultra-large B cells due to the computational complexity.

We also investigated the impact of sample size on the variability metrics using an external set of B cell ultra-large datasets with more than 10k cells (10X droplet-based) [17]. The 10k B cell data was compared to the B cell (naïve B cell; 50 cells) in TMS droplet-based data (see Methods). Overall, SD, IQR, scran, Seurat and BASiCS showed a reduction in the variability estimated for the 10k B cells compared to the TMS B cell dataset. Surprisingly, CV, FF, DESeq2, DM and edgeR showed increased gene expression variability for the 10k B cells. The higher average gene expression in the 10k B cells (reflective of expression captured for a higher number of cells) may explain the performance of the genes that showed increased expression variability, as these metrics largely relied on the mean expression during estimation. This result highlights how the same cell type sequenced by the same method but with different sample sizes can impact the estimation of gene expression variability (Figure 2b). This analysis identified scran and Seurat as having better performance for this criterion as they showed the least amount of change for the two sample sizes tested.

### Investigating the impact of different data structures and biological properties on metric performance

Measuring biological variability is challenging because of how scRNA-seq data is structured. For example, an excess of zeros, low average expression, and variability that is influenced by both technical and biological sources can be difficult to untangle, and these aspects all result in an increased amount of noise in scRNA-seq data compared to bulk level data. Therefore, it is important to select the metric that measures ‘true’ biological variability and is not driven by the noise that can be sourced back to these specific data structures alone.

Our study investigated how different data structures influence the performance of the 14 metrics (Figure 3, Supplementary figures 1 & 2) and found that scran and BASiCS performed well in most of the comparisons of the TMS cell type data. We explored the influence of features like the number of zeros a gene has for all cells, the mean expression value and the gene length. Although each metric handles zeros differently, we can still observe the increase in noise coming from the low signal genes in all the comparisons (Supplementary figure 2a). Additionally, most of the metrics preserved the mean-variance relationship except LCV, where lower average expression tends to correlate with a higher dispersion value (Supplementary figure 2b). Therefore, it is beneficial to pre-filter lowly expressed genes prior to any downstream analysis. Interestingly, even though the regression-based metrics and BASiCS considered the mean-variance relationship, the impact of data structures in these metrics showed diverse patterns. Gene length did not seem to impact the metrics when estimating gene expression variability (Supplementary figure 2c).

**Figure 3:**
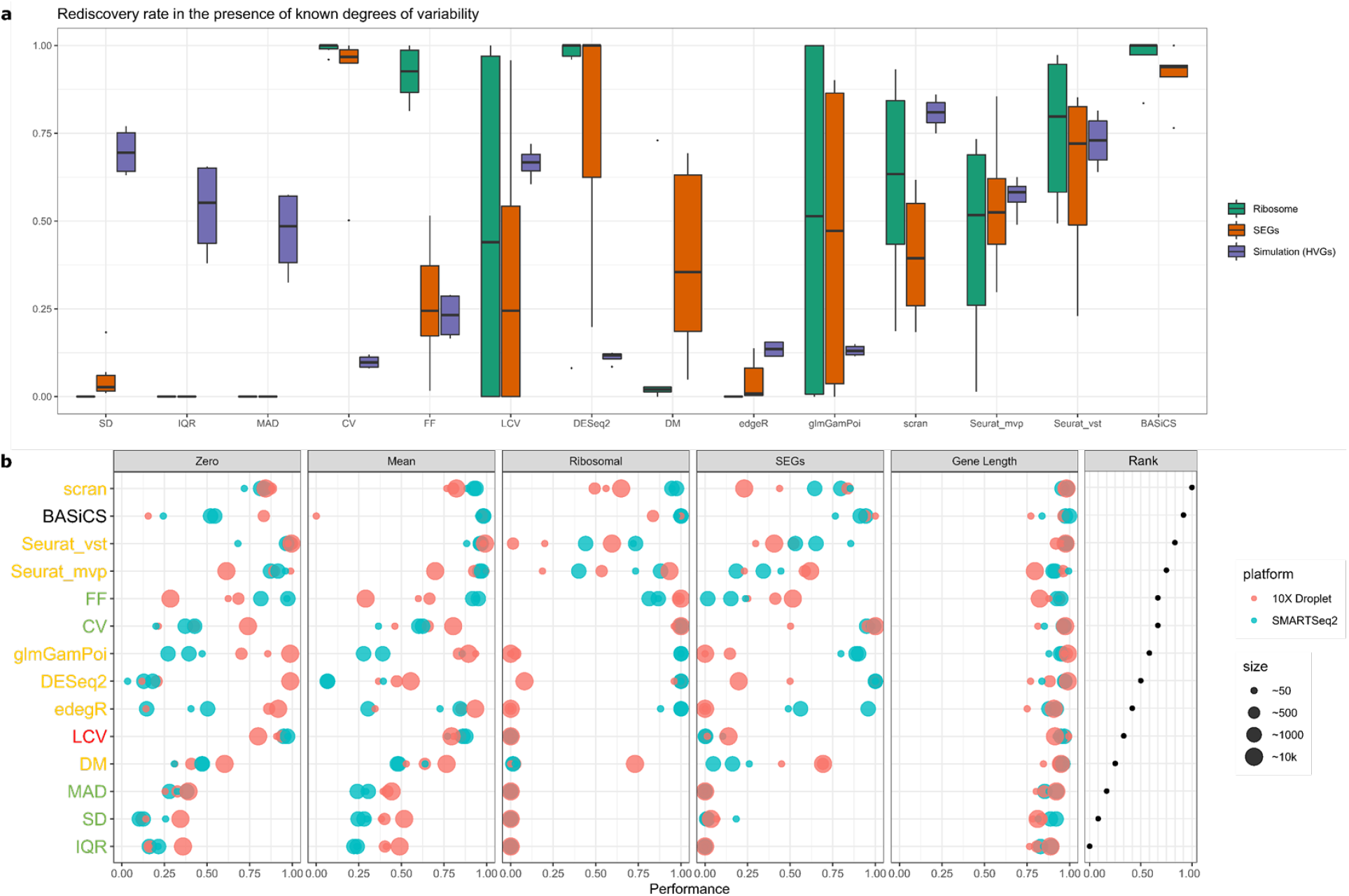
Metric performance in the presence of known degrees of variability and overall ranks for each metric. a) Boxplots for the rediscovery rate for each metric based on ribosomal genes, SEGs and simulations (HVGs). DM and BASiCS were not included as missing gene length and batch information, respectively. b) Bubble plot of metrics evaluation score across three cell types with two sequencing platforms. Each dot represented a dataset with its indicated number of cells. Each row represents a metric coloured by one of the four metric categories. Each section represents one main criterion, including proportions of zero per gene (zero), mean expression (mean), ribosomal genes (ribosomal), stably expressed genes (SEGs) and gene length. The higher value indicates a stronger performance score whereas 1 indicates perfect performance.

### Investigating the impact of known degrees of variability on metric performance

To investigate the performance of the variability metrics under controlled settings, we generated two types of control datasets to include in our evaluation. Firstly, a negative control dataset was used where no biological variability was measured [18]. We measured the overall estimated variability distribution dispersion from each metric for this dataset. A lower dispersion value demonstrated the capability of a metric to detect biological variability rather than the total variability. As a result (Supplementary figure 3), most of the basic statistics metrics performed well except for CV. The ranked-based approach for LCV could not be used for this comparison because the final estimated values always ranged from 1 to 100. Among the GLM-based metrics, scran performed best. Secondly, a set of four simulation studies under different parameters were used, where 200 genes were known to be highly variable (see Methods). Metrics were assessed based on the rediscovery rate (Figure 3a), and scran, Seurat_vst and SD obtained on average 70% of the rediscovery rate while DM, edgeR and DESeq2 only achieved around 20%.

We also evaluated the metric performance on biologically-relevant gene sets like ribosomal genes and stably expressed genes (SEGs) [19], where there is an overall expectation that these genes will have more stable gene expression. The gene rediscovery rate reflects how many of these genes’ expression variability are ranked within the first quartile of the metric’s gene expression variability distribution, indicating relatively stable expression (see Methods). The generic statistics metrics generally performed poorly by having a low rediscovery rate (Figure 3a and Supplementary figure 2d & 2e), however, CV was outstanding in its preservation of these biologically relevant genes. LCV showed very poor estimation of variability even though it performed well on eliminating the dependency from data structures. Most of the metrics based on regression model metrics performed well in preserving the SEGs, except for edgeR. Notably, DM excluded most of the ribosomal genes and some SEGs within its internal pre-filtering step. BASiCS performed consistently well in measuring the biologically-relevant SEGs, especially for the dataset collected from the droplet-based sequencing method. Taking into account the performance results based on both sets of control datasets and biological-relevant gene sets, scran was the best performing metric under these scenarios.

Both B cells and endothelial cells are types of abundant cells that can exist in multiple tissues, and we used this property to gain further insight into the performance of the best-performing metric, i.e., scran. Only tissues with more than 70 cells were included, resulting in 5 tissues for B cells and 7 tissues for endothelial cells. We hypothesised that the HVGs that were commonly detected between different tissues would be important for supporting cell type-specific roles while the HVGs unique to a tissue would support tissue-specific effects. To test whether scran was able to make this distinction, we selected the top 500 HVGs from scran and compared them to the top 500 HVGs from CV for each tissue-cell-type combination. As shown in Supplementary Figure 4, the HVGs measured using scran showed much more significant overlap across tissues compared to CV. The HVG gene lists measured from CV showed greater similarities to a random gene list as both gene lists show non-significant overlaps among tissue origins, suggesting a lack of true signal. Additionally, we confirmed that the HVGs measured by scran that were commonly detected between different tissues included many important cell type-specific markers (*Cd19, Cd83, Cd24a, Cd72* and *Cd48* for B cells; *Cd9, Tmem66* and *Tmem204* for endothelial cells). This analysis further indicates scran’s strength in performance at estimating gene expression variability.

### Fluctuations in cell-to-cell variability help explain the complex B lymphocyte differentiation process

Our comparative investigation into metric performance for estimating gene expression variability identified scran as the metric with the strongest overall performance. Hence, this metric was next applied to investigate how variability in cell-to-cell expression levels changed in HSC and B lymphocyte lineages and to identify specific markers of the differentiation process. We used the gene expression data from FACS Smartseq2 TMS marrow tissue and only incorporated cell types that had at least 100 cells in either young or old age groups (Supplementary Figure 5). The cell types included were late progenitor-B, precursor B, immature B and naïve B cells (Figure 4a). Gene expression levels were adjusted for the age effect using a regression model, given that the precursor B cells showed an overall significant difference between young and old groups (Supplementary Figure 6).

**Figure 4:**
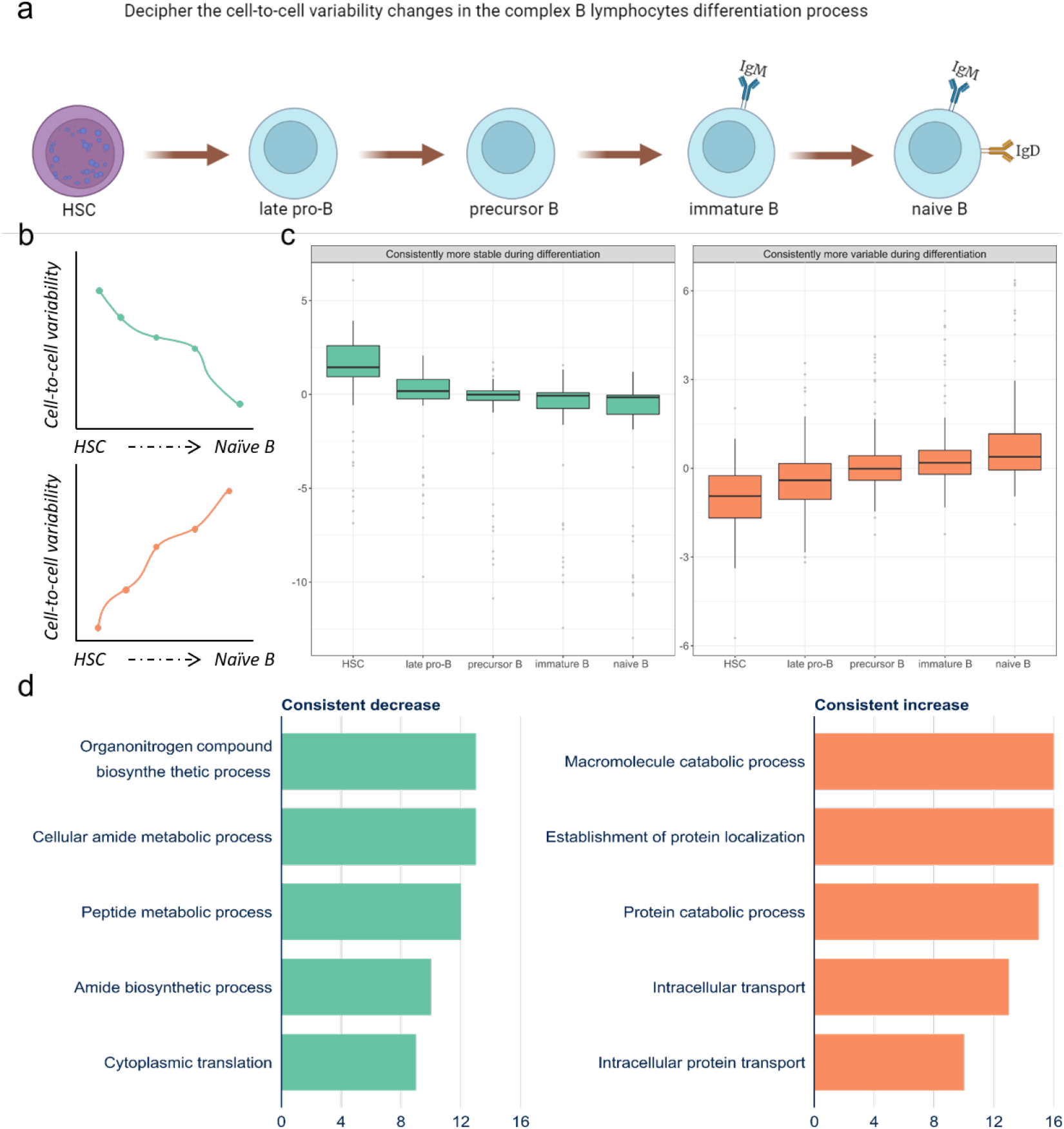
Deciphering the cell-to-cell variability of B lymphocytes during differentiation from TMS data. a) Workflow illustration of the variability estimation from HSC to multiple B lymphocytes. b) Line plot illustrating the two patterns of variability under study – increased stable and increased variable genes consistently during the differentiation process. c) Box plots showed the estimated variability for identified genes for different cell types. d) Barplots showed the top 5 significant pathways related to the consistent decrease and increase genes with GO biological process databases. Green represents consistently stable, while orange represents consistently variable genes.

Cell proliferation normally occurs through cell division that is highly associated with the G2M phase in the cell cycle. As expected, we saw that most of the cells in HSC and progenitor B cells were in G2M and S phase based on a list of markers [20] where they were prepared to proliferate while the cells in other cell types remained in G1 phase (Supplementary figure 7). In general, B lymphocyte differentiation occurs in the bone marrow from HSC through a complex transcriptional process, resulting in various B cell types [21]. Therefore, we re-constructed the lineage trajectory with a preset start [22] at HSC and observed the major trajectory from late progenitor-B, precursor B, immature B to naïve B cells (Supplementary Figure 8), which aligned with the underlying biology. TMS marrow data supported the differentiation process because the major trajectory reconstruction showed transitions from HSC, late progenitor-B, precursor B, immature B to naïve B cells.

Identifying genes with changes in gene expression variability that show consistent trends during the lineage transition may be an insightful way to identify new markers or regulators. For example, a gene that is consistently decreasing its expression variability may reflect a gene whose expression is increasingly stable as it transitions from HSC to naive B cells. On the other hand, genes that have consistently increasing expression variability may be related to cell-type-specific states during the differentiation processes [23].

We assessed the cell-to-cell variability changes along with the B lymphocyte lineages by identifying two gene expression patterns that were consistently more variable or stable in the differentiation process (Figure 4b). Each pattern consisted of 89 consistently variable genes as well as 47 consistently stable genes (Figure 4c). On average, there was an increased number of variable genes along the differentiation process relative to the stable genes. The top five markers for two patterns were *Ccr7, Tnfrsf13c, Cd19, Grap* and *Itm2b* for the consistently variable genes, as well as *Oaz1, Rpl31, Rpl38, Rpl28* and *Ccl9* for the consistently stable genes (Supplementary figure 9a and 9b).

Three cell differential markers, *Ccr7, Tnfrsf13c, Cd19*, showed the greatest variability changes along the differentiation process. Studies showed that these markers play essential roles in regulating immune-cell trafficking [24], B cell survival [25] and B cell developmental process [26]. Furthermore, *Grap* helps *Erk MAP kinase* activation by connecting the B cell antigen receptor, which provides the signal communications between a surface receptor to the DNA in the nucleus [27, 28], whereas *Itm2b* is known as a target of B-cell lymphoma 6 protein repression [29]. The genes that showed increased variability during cell differentiation belonged to pathways that were involved in B cell maturation. Conversely, most of the genes that were consistently more stable include the housekeeping gene *Oaz1* and ribosomal genes (*Rpl31, Rpl38*, and *Rpl28*). These genes are typically required for the maintenance of basic cellular functions.

The consistently variable genes showed more lymphoid-related specificity based on the percentage of expressed cells while consistently stable genes showed more generalised expression across different cell types (Supplementary Figure 9). Additionally, we performed pathway overrepresentation analysis of the two groups of genes to investigate their potential roles in B cell differentiation. Pathway analysis of increased variability genes showed a strong relationship with nitric oxide in the immune response whereas stably expressed genes most strongly associated with ribosome and peptide metabolic process (Figure 4d, Supplementary Table 3).

### Investigating cell-to-cell variability changes during ageing in HSC and naïve B cells

To decipher transcriptional heterogeneity changes in ageing, we measured the cell-to-cell variability for HSC and naïve B cells in the young and old groups, respectively (Figure 5a) using scran. We used these datasets to investigate how ageing impacts the variability of gene expression in these two functionally-relevant cell types, HSCs and naïve B cells. Overall, there was a slight decrease in variability observed in old HSC compared to young HSC (Wilcoxon test, marginal P-value = 0.04, Supplementary Figure 10a) and no significant effect in naïve B cells (Wilcoxon test, marginal P-value = 0.7, Supplementary Figure 10a). This result aligns with the expectation of stem cell exhaustion and a decline in B lymphopoiesis with ageing [30]. Genes that were highly variable in the young group versus the old group were more prevalent in both cell types (369 in HSCs, 327 in naive B cells). In contrast, there were 220 genes for the HSC and 269 genes for naïve B cells that were more variable in the old group versus the young group. Most of the enriched pathways were associated with the genes that reduced cell-to-cell variability in both cell types. Some shared pathways like ribosome and COVID-19 pathways may reflect the fluctuations occurring in tightened protein synthesis in ageing. In addition, DNA replication and cell cycle pathways are specifically enriched for the ageing HSC, which may explain the reduced activation of proliferation from a quiescence state to sustain hematopoiesis [31]. On the other hand, pathways like oxidative phosphorylation that are enriched for the aging of naïve B cells may present the loss of functions for cell energy and proliferation [32] (Supplementary figure 10b and 10c).

**Figure 5:**
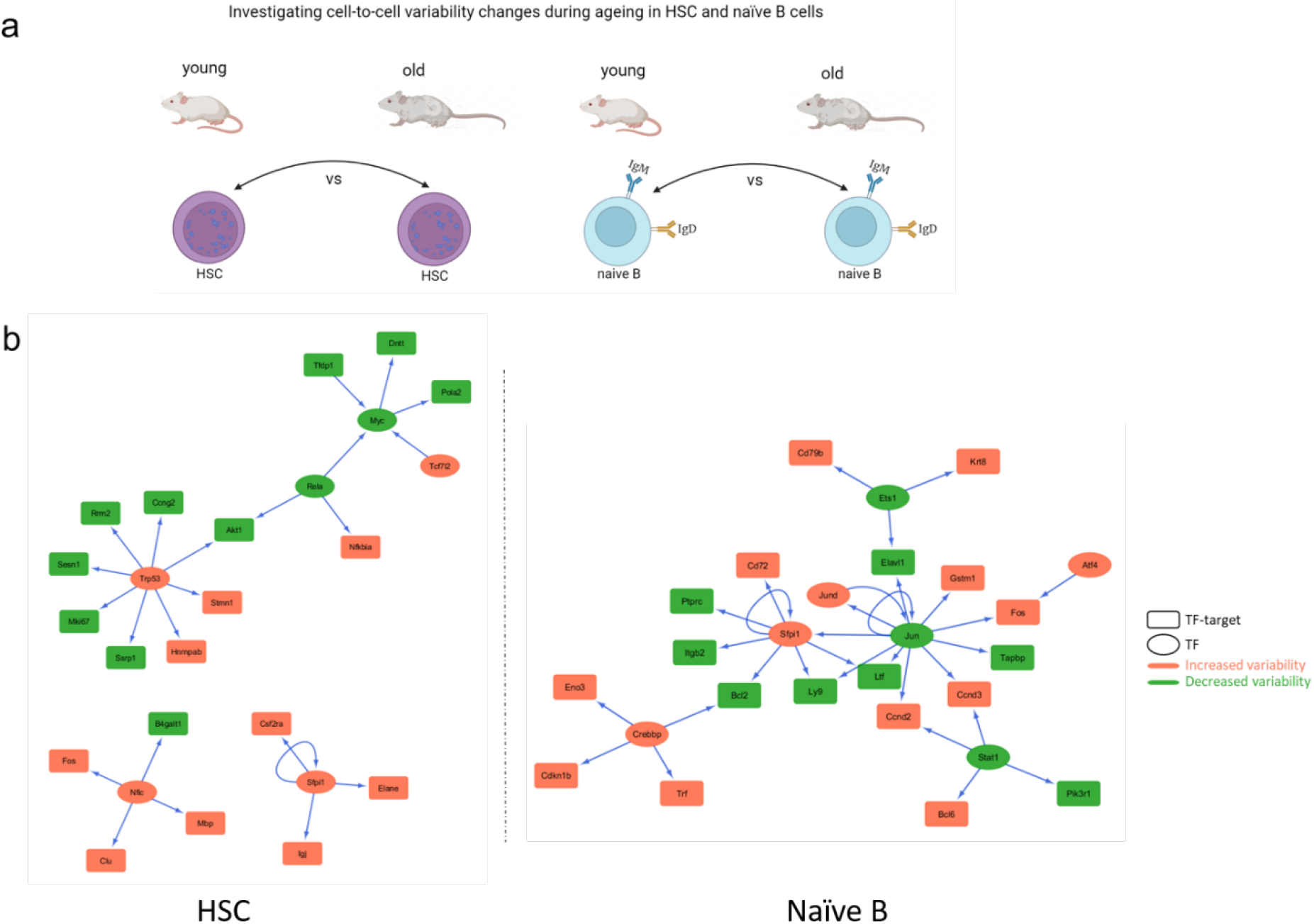
Deciphering the cell-to-cell variability of ageing in HSCs and naïve B cells from TMS data. a) Workflow illustration of the differentially variable analysis between young and old HSC and naïve B cells. b) TF networks were identified for HSC and naïve B cells based on the differentially variable genes. Square represented a TF while an oval represented its target. Green represents decreased variability in the old compared to the young group and red represents increased variability.

The development and proliferation of progenitor cells into lymphoid and myeloid cell lineages are tightly controlled by transcription regulation networks [21]. To assess alterations of the transcriptional regulation, we constructed regulatory networks based on the genes that were significantly differentially variable between young and old groups, which have been identified as transcription factors (TFs) and their known TF targets [33]. The functions of the networks were highly associated with stem cell differentiation and cell maintenance (Supplementary figure 10d). Interestingly, the analyses resulted in one connected network in naïve B cells and three smaller non-connected networks in HSC. Most DV genes that act as TFs have shown an increase in gene expression variability in older cells which may explain the dysregulation of regulatory mechanisms in cell-type-specific ageing. Additionally, we identified a shared TF, *Sfpi1*, which demonstrated consistently increased variability in ageing for both cell types (Supplementary figure 10e). However, the targets associated with *Sfpi1* presented unique expression changes for HSC and naïve B cells. *Sfpi1* is known as one of the primary lineage determinants that guide the multi-potential progenitors to establish a low-level expression of a mixed lineage pattern of gene expression, therefore favoured in lineage-specificity [34]. Increasing variable *Sfpi1* expression led to more variable targets’ expression such as *Csf2ra, Elane* and *Igj* in HSC while resulting in more stable expression of targets like *Bcl2, Ltf, Ly9, Itgb2* and *Ptprc* in naïve B cells.

## Discussion

Measures of cell-to-cell expression variability have been used extensively in the analysis of scRNA-seq data to understand heterogeneity and are often included in feature selection and dimension reduction steps. However, the applications of cell-to-cell variability measurements should not be restricted to selecting highly variable genes but incorporating the investigations of gene expression variability changes within and between conditions. Subtle changes in cell-to-cell variability for a single gene may be helpful for detecting shifts in gene expression heterogeneity, especially during differentiation processes or ageing. Studying cell-to-cell variability at the cell type level provides an interpretation of how gene expression variability is influencing regulation more directly than without acknowledging cell type groupings or modelling at the bulk level. There are tools such as MDSeq that use a novel re-parametrisation of the negative binomial to provide flexible GLMs on gene expression variability analysis, and scDD that identify the distribution changes under four scenarios [35, 36]. However, these proposed methods focus on detecting the changes between conditions and not for estimating a single condition.

One study recently highlighted the need to account for variability to accurately determine differentially-expressed genes [37]. Failing to accurately account for such variability may lead to spurious biomarker identification that does not truly explain the underlying biology. But one of the greatest challenges in single-cell analysis is determining the best way to compute measures of variability. Here, we conducted a systematic evaluation of the metrics for gene expression variability and concluded that no metric performed consistently well in all criteria. Different classes of metrics had stronger performance with respect to specific criteria and this is likely to be a result of the limitations of the underlying algorithm. Our benchmarking analysis demonstrated that the widely-used metric CV might introduce artificial variations due to the low mean expression resulting from excessive zeros in scRNA-seq data. In fact, our results demonstrated that most of the regression-model-based methods outperformed other metric categories which suggests that accounting for mean dependency in measuring gene expression variability is important.

Our study identified markers that had variable expression in the HSC to B lymphocytes differentiation process in bone marrow data. These types of variation changes that are continuous and subtle may mimic the continuous proliferation and differentiation processes for stem cells. Modelling consistent trends in increasing gene expression variability during the B lymphoid differentiation process identified not only genes that were known B cell maturation markers but also markers that control communication of signals through B cell antigen receptor connections. These consistently increasing expression genes represented the active cell fate decisions along with the differentiation, which is vitally important to autoimmunity and immunodeficiency [38]. Conversely, we identified *Oaz1*, which is known as a negative regulator of cell proliferation, together with other known ribosomal housekeeping genes that showed the most significant patterns in decreasing variability as cells transition from HSCs to B lymphocytes.

The role of transcription factors in modulating cell-to-cell variability can be useful for understanding how heterogeneity affects regulatory networks in aging. We identified genes that were differentially-variable between old and young mice, and we used these genes as input to construct regulatory networks based on the TFs for HSC and naïve B cells. Our results identified a shared TF *Sfpi1* that was consistently variable for both HSC and naïve B cells networks has been identified as a key dosage-dependent regulator of several hematopoietic cell lineage development and is associated with cell fate decisions [39].

Interestingly, *Sfpi1* also targeted a more variable *Cd72* in aged naïve B cells, in which *Cd72* is a primary regulator that appears to mediate B-cell and T-cell interaction. Moreover, differentially varied *Sfpi1* targets were distinct for HSC and naive B cells and their expression changes direction also differed. The cell-type-specific regulatory networks from our studies identified the unique TF-target interplay changes during ageing process, which may provide novel targets to predict the consequences of ageing HSC and B cells.

There are multiple reports of increased cell-to-cell variability with ageing in different cell types. However, some studies have also reported uncertain or inconsistent levels of cell-to-cell variability changes in ageing by similar experimental settings. One reason could be the biological differences in ageing processes among different species, organs and conditions. However, it is also possible that the metrics are misused in estimating gene expression variability. With an increasing prevalence of multi-source and multi-condition datasets, the underlying data structures should be carefully considered before applying any metric to measure the gene expression variability. Overall, our analysis provides evidence in applying model-based metrics like scran to accurately estimate biologically relevant gene expression variability, where we demonstrate its ability to provide novel insight into complex conditions, like differentiation and ageing.

## Conclusions

Cell-to-cell variability is important for understanding cellular heterogeneity under different biological conditions. Here, we have performed a comprehensive benchmarking study to explore the performance of 14 metrics that quantifies cell-to-cell variability. Based on the evaluation, we found that the gene expression variability estimates from the scran R package have the best performance in recapitulating the biological cell-to-cell variability and are independent of the data structures. We show the significant impact of different levels of cell-to-cell variability under two important biological processes such as differentiation and ageing. Our analyses demonstrate that cell-to-cell variability changes reveal critical roles in not only maintaining cell functions but also accurately capturing the key regulators. We conclude that the cell-to-cell variability should not primarily be considered as noise in analysing scRNA-seq data, but also provide remarkable information in understanding complex biological processes.

## Methods

### Overview of the metrics

The 14 metrics used in this study are commonly used to quantify gene expression variability and represent four distinct categories: generic metrics, local normalisation metrics, regression-based metrics and Bayesian-based metrics (Figure 6). These metrics are either generic or specifically designed in a package for analysing transcriptomic data. We used log CPM-normalised gene expression as input for the generic metrics and local normalisation metrics categories, and raw gene count matrix for the metric in other categories. One of these metrics, LCV, was originally designed for bulk RNA-seq data so we have modified it to be compatible with assumptions for scRNA-seq data. Unless stated, the metrics were run with default parameters as described in their respective vignettes and papers.

**Figure 6:**
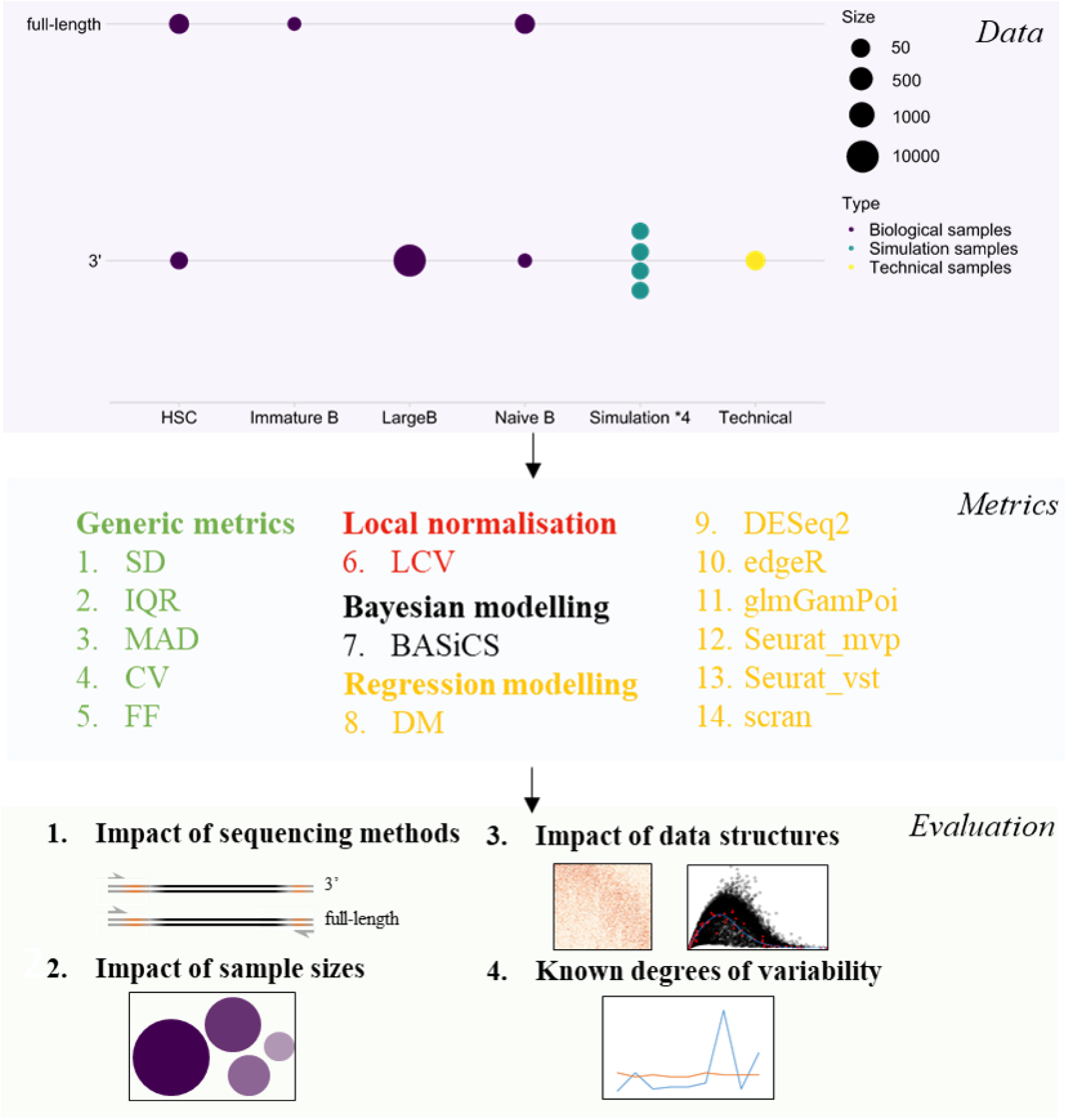
Schematic summaries of the benchmark workflow to evaluate the performance of cell-to-cell variability metric for scRNA-seq data. A panel of 14 metrics that perform the estimation of cell-to-cell variability were evaluated on both experimentally-derived and simulation data from two main sequencing platforms with different sample sizes. Evaluation includes the impact of sequencing platforms, sample sizes, data structures and the recapitulation of known degrees of variability. Other than generic metrics, other metrics are designed for analysing transcriptomic data and implemented in the corresponding packages.

### Generic metrics

The five metrics in this category include median absolute deviation (MAD), interquartile range (IQR), standard deviation (SD), coefficient of variation (CV) and the Fano Factor (FF). SD, MAD and IQR have been interpreted as a stochastic disturbance in the data which represents the uncertainty around the mean value, and have previously been applied to gene expression analyses [40]. Additionally, gene expression variability is known to show some dependence on the mean expression level, and the metrics CV (σ^2^/μ) and FF (σ/μ) are two of the most commonly-used metrics for modelling variability in scRNA-seq data that explicitly acknowledge a mean-variance dependency [41, 42].

### Local normalisation metrics

The Local Coefficient of Variation (**LCV**) [43] was first developed to rank the expression variability of each gene relative to genes with similar local expression values in bulk RNA-seq data. In our adaptation to scRNA-seq data, we have tried to retain as much of the ranking algorithm and the relationship between the mean and CV. LCV starts with ordering each gene according to its mean expression and then assigns its corresponding CV into the user-defined width. It re-calculates the local CV as the quantiles of the CV within the window for each gene. In such a way, this metric rescales variation within the whole gene population to a user-defined range.

### Regression-model based metrics

Novel model-based approaches have been introduced to regress unwanted technical variation, to improve the accuracy and detection of ‘true’ biological variation in the gene expression data. Several metrics in this category were included in our comparative study because of the different model assumptions that they adopt.

#### DESeq2

[44] estimates dispersion by applying a generalised linear model of the form can be summarised as:

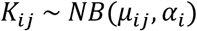

where the raw count matrix *K*_*ij*_ for gene *i*, sample *j* are modelled by a negative binomial distribution with fitted mean *μ*_*ij*_ and a gene-specific dispersion parameter *α*_*i*_. *α*_*i*_ reflects the relationship between the normalised mean expression and the variance for the observed gene *i* expression which is assumed to follow a log-normal prior distribution. Next, a curve is fitted to all gene-wise dispersions and further shrunk towards the expected dispersion value. The final dispersion estimates were used to represent gene expression variability.

#### DM

(distance to the median) Kolodziejczyk and Kim et al., [45] developed a novel method to account for the confounding factor of expression levels when calculating cell-to-cell variability. The distance between the CV^2^ of each gene with a running median was calculated and then corrected for using the gene length to eliminate the confounding effect. They refer to this expression-level normalised measure of gene expression heterogeneity as DM.

#### edgeR

[46] also models the gene counts using a negative binomial distribution and the estimation of the dispersion parameter was done in two steps. First, a common dispersion is estimated by maximising the adjusted likelihood function on the squared biological coefficient of variation (BCV). These common dispersion estimates are further generalised by binning and weighting the subsets of the dispersion grid, so-called trended dispersion. Second, the weighted likelihood empirical Bayes is applied to squeeze the tagwise (gene-wise) dispersions towards a common dispersion by obtaining posterior dispersion estimates. These tagwise dispersions were used to estimate the gene expression variability in this comparative study.

#### glmGamPoi

[47] leverages the inferior transformation approach, which outperforms DESeq2 and edgeR by substantially higher speed and the inference to be more suitable for scRNA-seq data by using efficient data representations. Through modelling under a Gamma-Poisson distribution, it also shows better estimates of the over-dispersion parameter on datasets with low counts.

#### scran

[48] performs a unique normalisation process by pooling cells to calculate multiple scaling factors, that leads to a more accurate estimate that improves downstream analysis. The total variance in expression for each gene is fitted with log-normalised expression by a mean-variance trend. To obtain an estimate of biological variability for each gene, the decomposition step is applied to the total variance by subtracting the fitted value (technical variability). The biological variability estimates were used to estimate gene expression variability in this comparison.

#### Seurat

[49, 50] performs normalisation by dividing gene counts for each cell by library size and further multiplying by 10 000. It includes two main methods to examine gene expression variability. One way is to use a binning method (default bin=20) on the average expression and then calculate z-scores for dispersion within each bin for each gene (Seurat_mvp). This approach aims to control for the strong mean dependency when calculating the dispersion. Alternatively, it fits a trend line to the relationship of mean and variance using LOESS then standardises and scales the feature values using the observed mean and expected variance (Seurat_vst). Both methods will be analysed in the comparison separately.

### Bayesian modelling metrics

#### BASiCS

[51] is the first method that obtains estimates of biological variation using a Bayesian hierarchical model. The batch information was included for measuring technical variability in BASiCS. The biological variability is measured by a residual measure of variability given by the departures from a global mean/over-dispersion trend, with Regression = TRUE for both data types.

### Datasets and pre-processing used for metrics evaluation

To compare the performance of the different metrics, we focused on investigating changes in expression variability from the Tabula Muris Senis (TMS) data at the cell type level. TMS is a large publicly available mouse atlas that includes scRNA-seq data, generated from both FACS-sorted single cells and through droplet sequencing, across 24 organs with rich transcriptome information [11]. FACS data allow for higher sensitivity and less sparsity whereas droplet-based sequencing enables more cells to be analysed. We downloaded the raw gene expression count matrix (FACS-sorted Smart-seq2 and 10X Genomics droplet-based) from the figshare repository (https://doi.org/10.6084/m9.figshare.8273102.v3) and normalised these data using the Seurat package (version 3) following the descriptions given in the original paper [52]. For our metric evaluation, only 24-month male mice were included for the downstream analysis. We estimated cell-to-cell variation exclusively at the level of a single cell type.

Marrow tissue was selected for its multipotent capabilities and involvement in the immune response. We also selected this tissue for statistical reasons because marrow tissue had the largest number of cells that were sequenced. Out of the 23 cell types in the marrow tissue, several cell types were carefully chosen due to the data properties, as summarized in Supplementary Table 1. HSC and Naïve B cells were two of the most representative and abundant cell types that were sequenced in the FACS Smart-seq2 technology. Corresponding cell types that were sequenced by droplet-based technology were also included as a comparison for the metric performance. The immature B cells were selected because they had a relatively smaller sample size and this is an important contrasting comparison to include because the sample size is critical in measuring gene expression variability estimation sensitivity [53]. In addition, we applied these metrics on B cells and endothelial cells from all tissues available in TMS to evaluate the metric’s ability to detect genes that are responsible for functional maintenance across tissues.

Beyond TMS, two external datasets were further tested as controls to investigate the metric performance. The first dataset was sequenced from 92 synthetic spike-in RNA molecules from the External RNA Controls Consortium (ERCC) to understand technical variability [18]. In theory, there is no genuine biological variability derived from this experiment, so this dataset represents a limited level of variability compared to other gene expression data. The second dataset was included because it has a much larger sample size, containing more than 10k sorted CD19+ B cells from fresh PBMCs which is 10 times greater than the TMS datasets [17]. As expression variability is reduced with sample size, this comparison demonstrates how well each matrix can handle big data.

Although there were studies that identified stably expressed genes, it is challenging to identify the ground truth for highly variable genes with real data. Therefore, four independent simulation studies were generated from parameters that were comparable to the TMS dataset via the Bioconductor package Splatter [53]. Simulation 1 (Sim1) and Simulation 2 (Sim2) consisted of 200 genes by setting the BCV parameters to 3-fold and 4-fold of the original parameter to imitate the different levels of variability. Additionally, to accommodate extremely noisy and sparse scRNA-seq data, we modified the dropout.mid from 2 to 10 on Sim1 and Sim2, denoted as Sim3 and Sim4. For each simulated dataset, the total number of cells was set to 400 and the total number of genes was set to 1000, where 200 of the genes were known to be highly variable.

The details of the datasets used in this study were summarised in Figure 6, Supplementary Table 1 and online repository, including the functions to calculate cell-to-cell variability that were implemented in the non-generic packages and the corresponding publication.

### Metrics evaluation score

To assess the performance of each metric, we measured the associations between each metric and characteristics of the data, such as the proportions of zeros, the mean expression, and the gene length. As DM required a pre-filtering step, it resulted in fewer genes available to be tested in the analysis (a loss of about 40% of genes) which affected its evaluation of performance in our comparison. Additionally, the log-transformed mean expression was used for all metrics, except for edgeR, DESeq2 and scran as they have different internal normalisation approaches that resulted in different mean expression values. Available gene length was retrieved for 18,000 genes from the mm9 reference by the goseq package (version 1.44.0) [54]. The degree of association was assessed by Pearson correlation coefficient and the corresponding R-value was kept as the measurement score. The higher R-value represented a stronger association between the estimated variability and other data characteristics, which reflected a greater influence on performance due to the data structures. For assessing the metric performance on genes that were expected to have low expression variability, we used ribosomal genes and mouse stably expressed genes (SEGs) [19] because they have housekeeping roles that are critical to maintaining fundamental functions of the cell. We scored the metrics based on the number of genes from these categories (ribosomal genes or SEGs) that were in the first quartile of the estimated variability distribution for each metric. The higher proportion of genes that followed such criteria, the better performance of the metric because it accurately calculated the lowly variable genes. For the simulation data, the rediscovery rate was calculated to be determined by the number of identified HVG genes.

### TMS B lineages datasets and pre-processing

To understand how cell-to-cell variability changes in B cell development, we included the FACS-sequenced data from 3-month and 24-month old mice as they have a relatively higher abundance of sequenced cells in TMS compared to other time points. We selected the B lineage-related cell types that have at least 100 cells in either age group, resulting in HSCs, late progenitor-B, precursor B, immature B (IB) and naïve B cells (NB). Raw data was processed according to descriptions in the original paper for all cell types and conditions. To increase the sample size for analysis of the variability patterns in HSC and B cell lineages, we merged both age groups for each cell type to regress out the age effect using a design matrix. Conversely, 3-month and 24-month HSC and naïve B cells were used for measuring differentially variable genes in ageing.

### Trajectory analysis

Pseudotime trajectory analysis was performed to determine how cells transition from one state to another based on the TMS B cell lineage datasets. We constructed the trajectory with Monocle3, which presets the starting point as HSC [22]. Cell pseudo-time values were applied and coloured for each cell identity on the UMAP to indicate the potential trajectory along each developmental process.

### Network of transcriptional factors and target genes

RegNetwork [33] was used to retrieve the transcription factors and their target genes. TF networks were constructed using Cytoscape (v3.7.1) [55], where the arrows denote a link from a TF to a target gene. TFs were annotated with rectangles and targets were annotated with circles. The colour indicates whether a gene was more variable or less variable in the old compared to the young group.

### Differentially variable (DV) gene testing

To increase the power for calculating DV genes, we first pre-filtered the genes based on the proportions of zeros. To determine the differentially variable genes, we calculated the cell-to-cell variability (CCV) difference δ measured by scran, which is defined as δ = CCV (Old)-CCV (Young). Then δ was normalised as a z-score and further converted to p-values with the assumption that all z-scores follow a normal distribution. In this paper, we selected the features whose p-value <0.05.

### Pathway over-representation analysis

To investigate gene sets at the molecular and functional level, we performed enrichment analysis on GO biological processes pathways by using MsigDB [56]. The top 5 significant pathways were shown in the barplot and ordered by their overlapped gene set size.

### Statistical analysis and data availability

Statistical analysis was performed using R version 3.6 (https://www.r-project.org/). All the scripts and analysis pipeline are available from the Github page: (https://github.com/huiwenzh/cell-to-cell-variability-changes-in-ageing). Plots were generated through the ggplot2 version 3.3.2 package unless stated.

## Availability of data and materials

The data used in this analysis are all publicly available. All datasets are described in the “Methods” section and supplementary Table 1 with all links. Codes to reproduce the presented analyses and figures are available at https://github.com/huiwenzh/cell-to-cell-variability-changes-in-ageing. Statistical analysis was performed using R version 3.6 (https://www.r-project.org/). Plots were generated through the ggplot2 version 3.3.2 package.

## Competing interests

The authors declare no competing interests.

## Funding

Australian Research Council Future Fellowship (FT170100047) and a Georgina Sweet Award to J.C.M

## Authors’ contributions

H.Z., J.V. and J.C.M formulated the problem. H.Z. designed the evaluation framework and analysis with input from A.T.F and J.C.M. H.Z performed the B cell ageing analysis with input from A.T.F and J.C.M. H.Z., A.T.F and J.C.M interpreted the results with input from J.V. H.Z. wrote the manuscript with input from A.T.F, J.V. and J.C.M. All authors read and approved the final manuscript.

## Acknowledgements

A sincere thank you to Prof. Di Yu from The University of Queensland Diamantina Institute and A/Prof. Xiao Dong from The University of Minnesota for valuable comments and discussion.

## Notes

### Competing Interest Statement

The authors have declared no competing interest.

