## Supplementary materials for "Measuring cell-to-cell expression variability in single-cell RNA-sequencing data: a comparative analysis and applications to B cell ageing"

### **Supplementary – Tables**

Supplementary Table 1: Detailed summary of the data used in comparing metric performance.

| Dataset name | Platform | No. of cells | No. of genes | Cell type | Reference |
| --- | --- | --- | --- | --- | --- |
| TMS-HSC-droplet | 10X Droplet-base | 486 | 20138 | Hematopoietic precusor cell | [1] |
| TMS-NB-droplet | 10X Droplet-base | 49 | 20138 | Naïve B cell |  |
| TMS-HSC-FACs | Smart-seq2 | 1174 | 22899 | Hematopoetic stem cell |  |
| TMS-NB-FACs | Smart-seq2 | 1166 | 22899 | Naïve B cell |  |
| TMS-IB-FACs | Smart-seq2 | 44 | 22899 | Immature B cell |  |
| PBMC_B_droplet | 10X Droplet-base | 10085 | 15858 | CD19 B cell | [2] |
| Technical variability data | 10X Droplet-base | 1015 | 92 | ERCC spike-in |  |
| Simulation1 | Splatter | 400 | 1000 |  | [3] |
| Simulation2 | Splatter | 400 | 1000 |  |  |
| Simulation3 | Splatter | 400 | 1000 |  |  |
| Simulation4 | Splatter | 400 | 1000 |  |  |

Supplementary Table 2: Kolmogorov–Smirnov’s D statistic measured the distribution difference between two sequencing platforms for the same cell types as well as the difference between two cell types for the same sequencing platforms.

|  | sd | iqr | mad | CV | FF | edgeR | DESeq2  glm |
| --- | --- | --- | --- | --- | --- | --- | --- |
| HSC_facs vs HSC_droplet | 0.188 | 0.08 | 0.07 | 0.20 | 0.20 | 0.92 | 0.30 |
| HSC_facs vs NB_facs | 0.05 | 0.20 | 0.11 | 0.20 | 0.47 | 0.18 | 0.25 |
| NB_facs vs NB_droplet | 0.33 | 0.27 | 0.12 | 0.58 | 0.55 | 0.99 | 0.76 |
| HSC_droplet vs NB_droplet | 0.12 | 0.13 | 0.07 | 0.27 | 0.33 | 0.24 | 0.39 |
|  | **DESeq2**  **glmgam** | **DM** | **LCV** | **Seurat_mvp** | **Seurat_vst** | **scran** | **BASiCS** |
| HSC_facs vs HSC_droplet | 0.70 | 0.04 | 0.03 | 0.14 | 0.23 | 0.27 | 0.75 |
| HSC_facs vs NB_facs | 0.24 | 0.07 | 0.029 | 0.14 | 0.073 | 0.10 | 0.26 |
| NB_facs vs NB_droplet | 0.88 | 0.03 | 0.03 | 0.16 | 0.25 | 0.15 | 0.98 |
| HSC_droplet vs NB_droplet | 0.37 | 0.08 | 0.05 | 0.15 | 0.22 | 0.15 | 0.37 |

Supplementary Table 3: Top 5 over-represented pathways for the consistently variable and consistently stable genes along the B lymphocytes differentiation process against GO biological process database.

| **Gene category** | **Pathway name** | **Genes in the pathway** | **q-value** |
| --- | --- | --- | --- |
| Consistently variable genes (89) | GOBP_PROTEIN_CATABOLIC_PROCESS | *Laptm5, Gga1, Rab7, Vps36, Bnip3l, Ubl4a, Gabarapl2, Rack1, Ube3a, Rad12a, Ube3q, Rad23a,Usap19,Rnf144a, Ubr4* | 1.84 e^-5^ |
|  | GOBP_ESTABLISHMENT_OF_PROTEIN_LOCALIZA  IZATION | *Laptm5, Gga1, Rab7, Vps36, Bnip3l, Ubl4a, Gabarapl2, Rack1, Vsp29, Ift27, Ripor1, Srpr, Nxt1, Unc93b1, Arcn1, Hif1a* | 2.05 e^-4^ |
|  | GOBP_INTRACELLULAR_TRANSPORT | *Laptm5, Rab7a, Bnip3l, Dtx3l, Vsp29, Ift27, Ripor1, Srpr, Nxt1, Unc93b1, Arcn1, Hif1a, Atp2a2* | 2.39 e^-4^ |
|  | GOBP_INTRACELLULAR_PROTEIN_TRANSPORT | *Laptm5, Rab7a, Bnip3l, Dtx3l, Vsp29, Ift27, Ripor1, Srpr, Nxt1, Unc93b1* | 2.93 e^-4^ |
|  | GOBP_MACROMOLECULE_CATABOLIC_PROCESS | *Laptm5, Gga1, Rab7, Vps36, Bnip3l, Ubl4a, Gabarapl2, Rack1, Ube3a, Rad12a, Ube3q, Rad23a,Usap19,Rnf144a, Ubr4, Dnase1l3* | 2.93 e^-4^ |
| Consistently stable genes (47) | GOBP_CYTOPLASMIC_TRANSLATION | *Rpl15, Rpl22, Rpl24, Rpl28, Rpl29, Rpl31, Rpl36a, Rpl38, Rps17* | 4.94 e^-10^ |
|  | GOBP_PEPTIDE_METABOLIC_PROCESS | *Rpl15, Rpl22, Rpl24, Rpl28, Rpl29, Rpl31, Rpl36a, Rpl38, Rps17, Gstm1, Gstm7, Cpq* | 6.5 e^-7^ |
|  | GOBP_CELLULAR_AMIDE_METABOLIC_PROCESS | *Rpl15, Rpl22, Rpl24, Rpl28, Rpl29, Rpl31, Rpl36a, Rpl38, Rps17, Gstm1, Gstm7, Cp1, Sirt3* | 6.5 e^-7^ |
|  | GOBP_ORGANONITROGEN_COMPOUND_BIOSYNTHE  THETIC_PROCESS | *Rpl15, Rpl22, Rpl24, Rpl28, Rpl29, Rpl31, Rpl36a, Rpl38, Rps17,Lmf1, Dhodh, Sirt3, Oaz1* | 3.85 e^-5^ |
|  | GOBP_AMIDE_BIOSYNTHETIC_PROCESS | *Rpl15, Rpl22, Rpl24, Rpl28, Rpl29, Rpl31, Rpl36a, Rpl38, Rps17, Sirt3* | 3.85 e^-5^ |

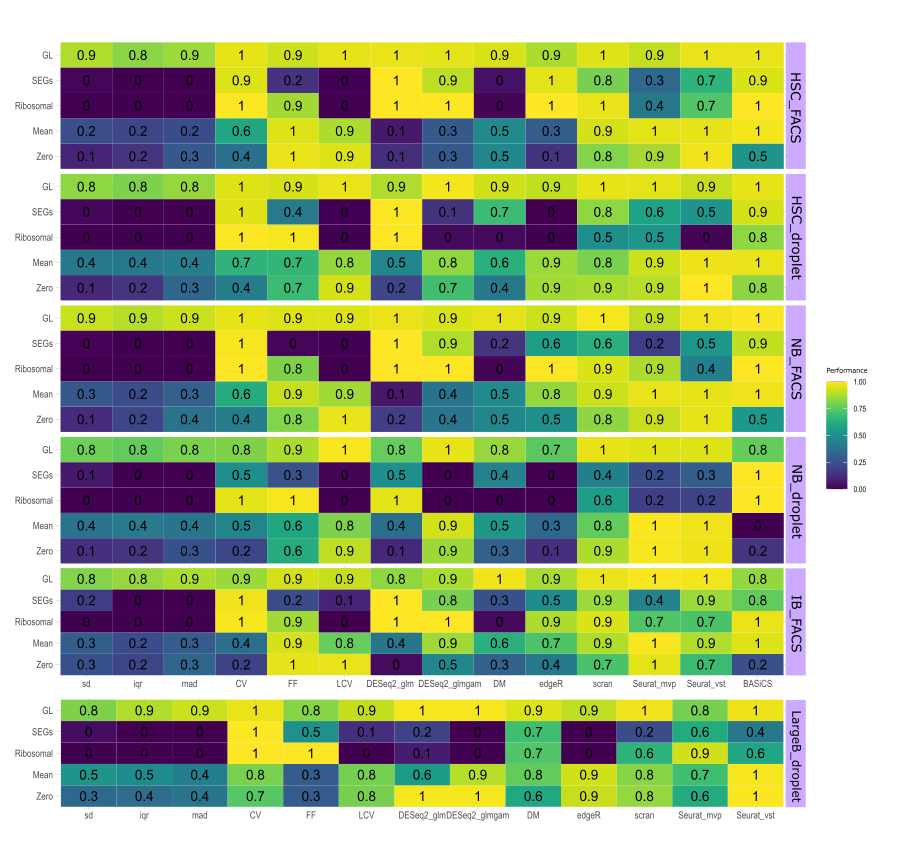

Supplementary figure 1. Heatmaps of metrics performance on three cell types with two sequencing platforms. Each role represented five main criteria which included proportions of zero per gene (zero), mean expression (mean), ribosomal genes (ribosomal), stably expressed genes (SEGs) and gene length (GL). Metric was highlighted by different categories. Data used for each evaluation heatmap wass labelled on the right, where the first 5 were from TMS datasets and the last one represented 10k B cell data. BASiCS was not applied to large B cells as lacking batch information and computational complexity.

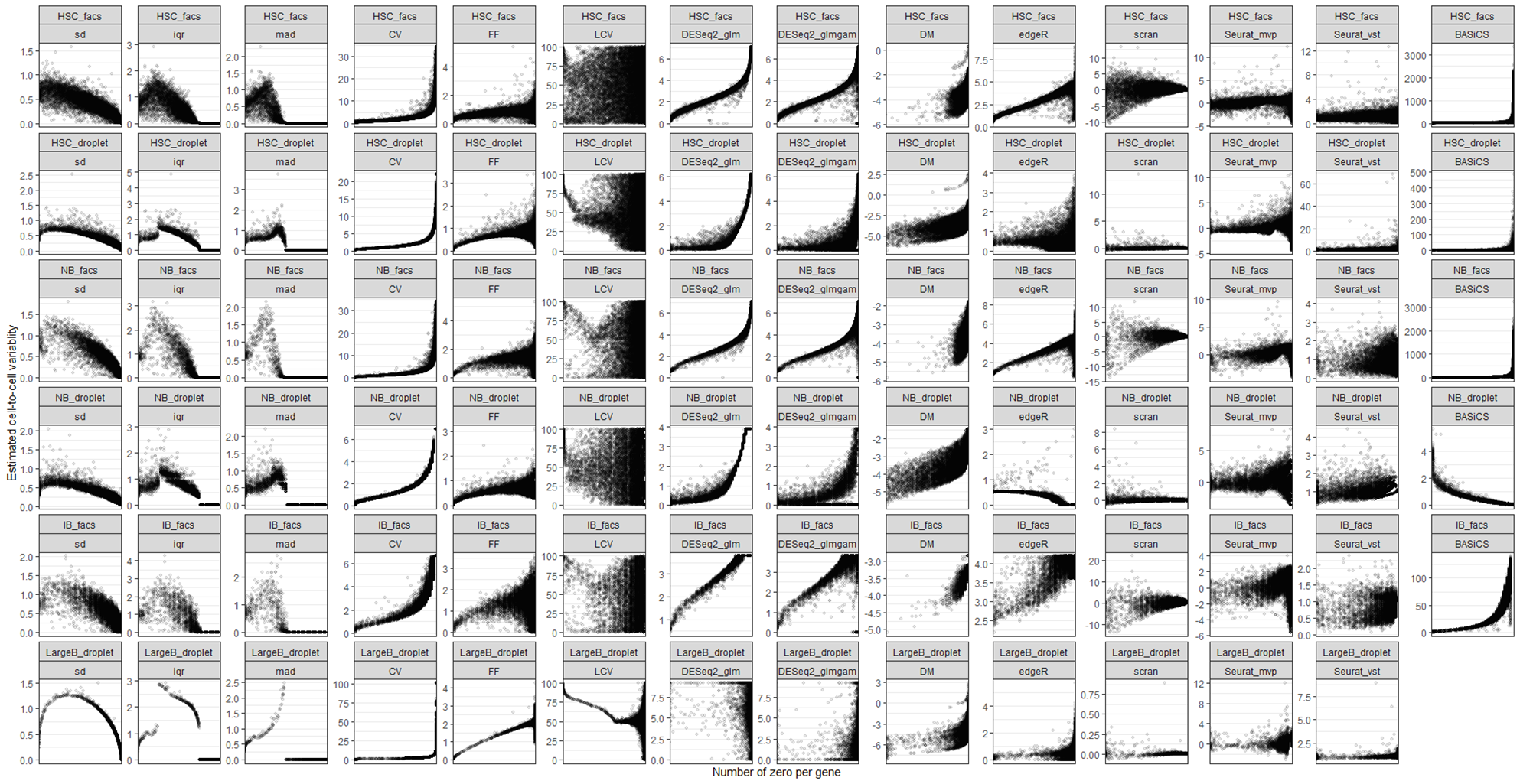
**A**

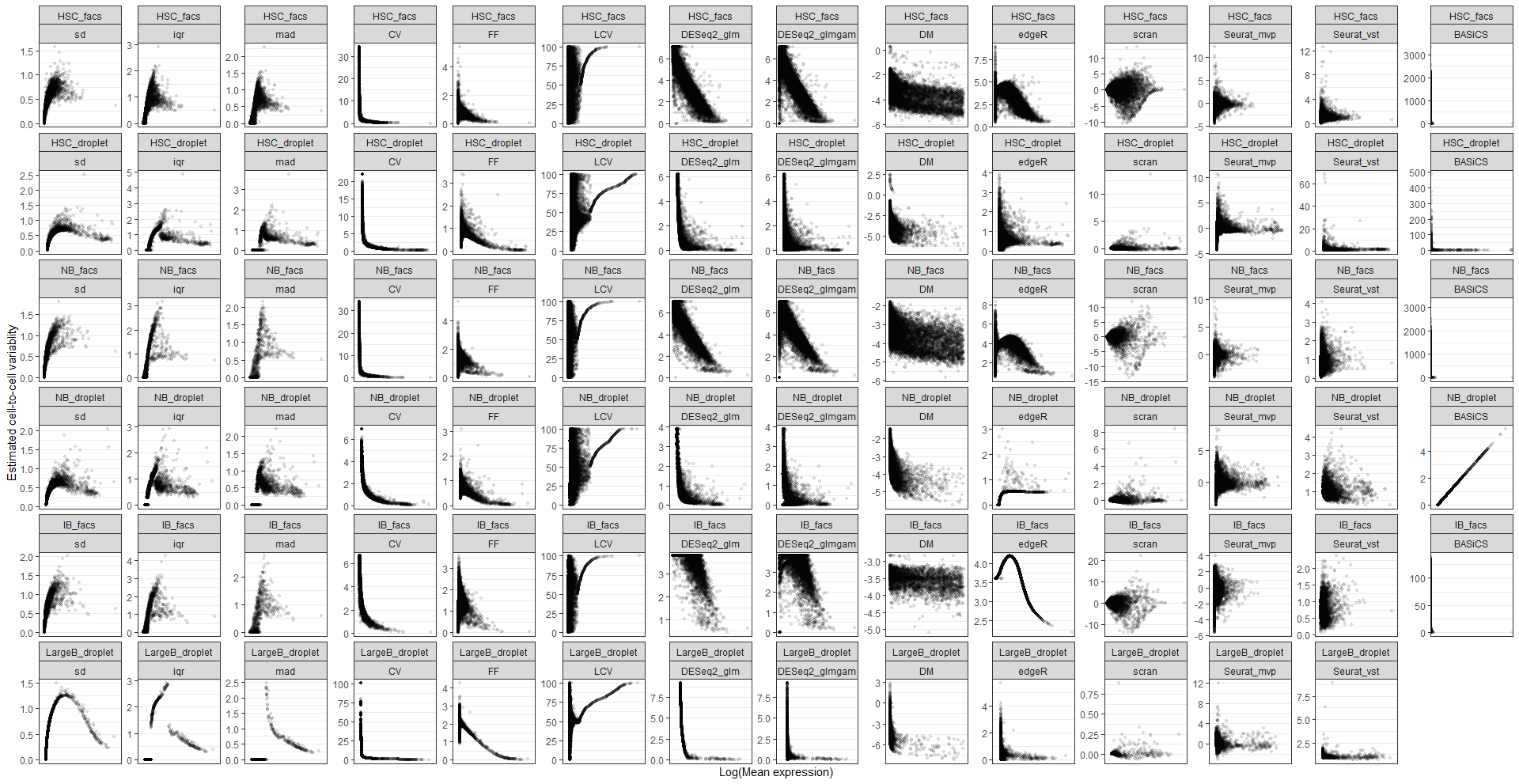
**B**

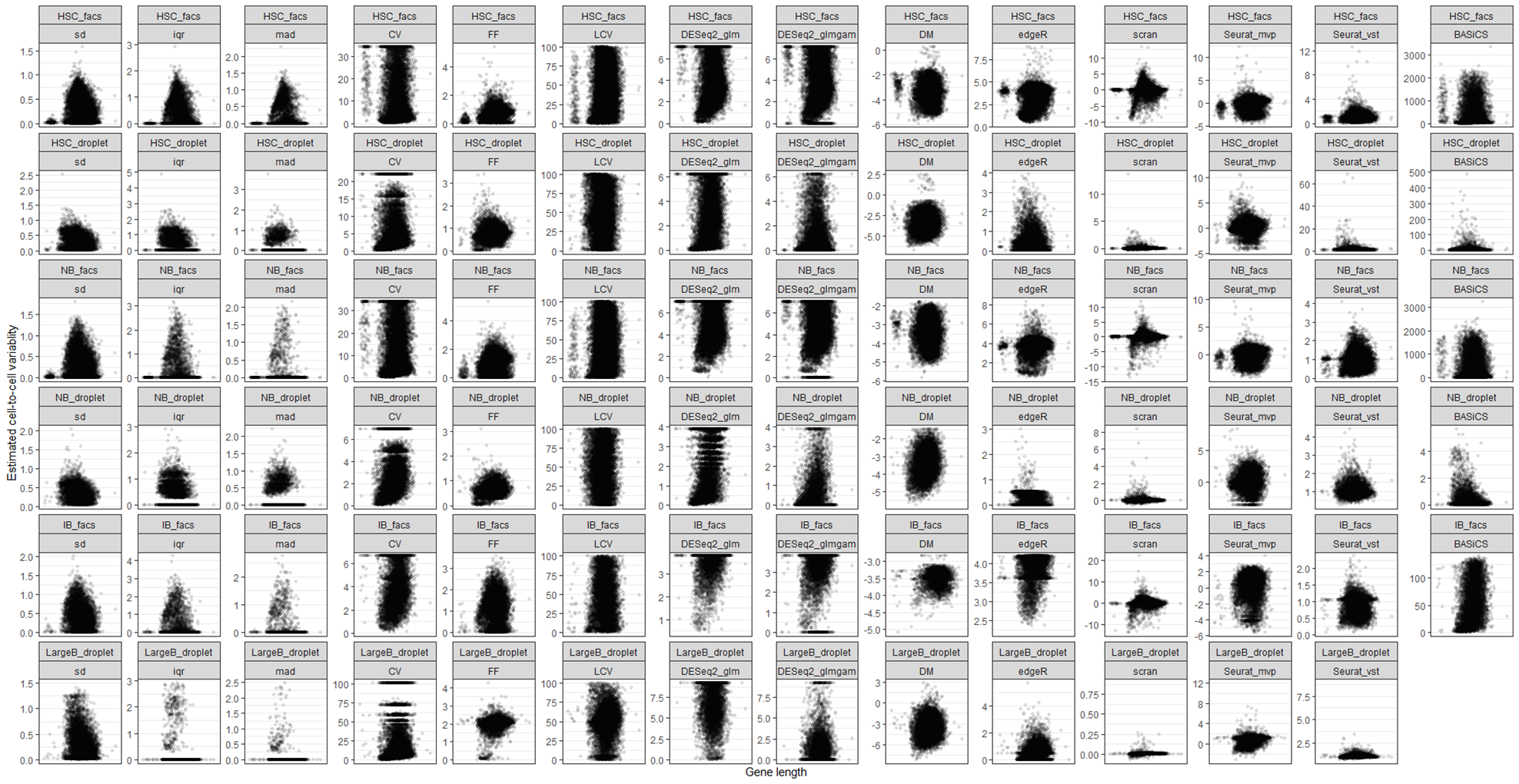
**C**

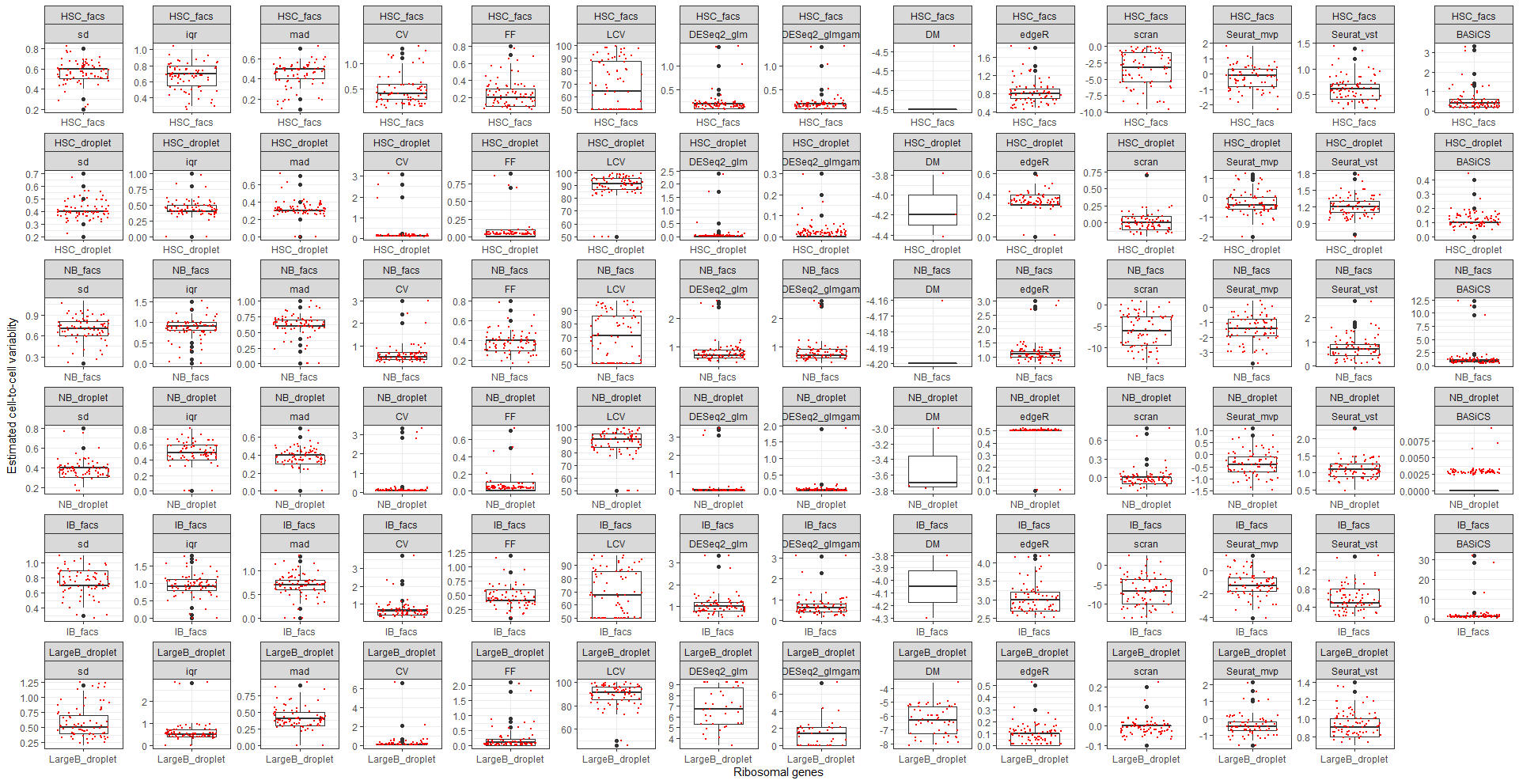
**D**

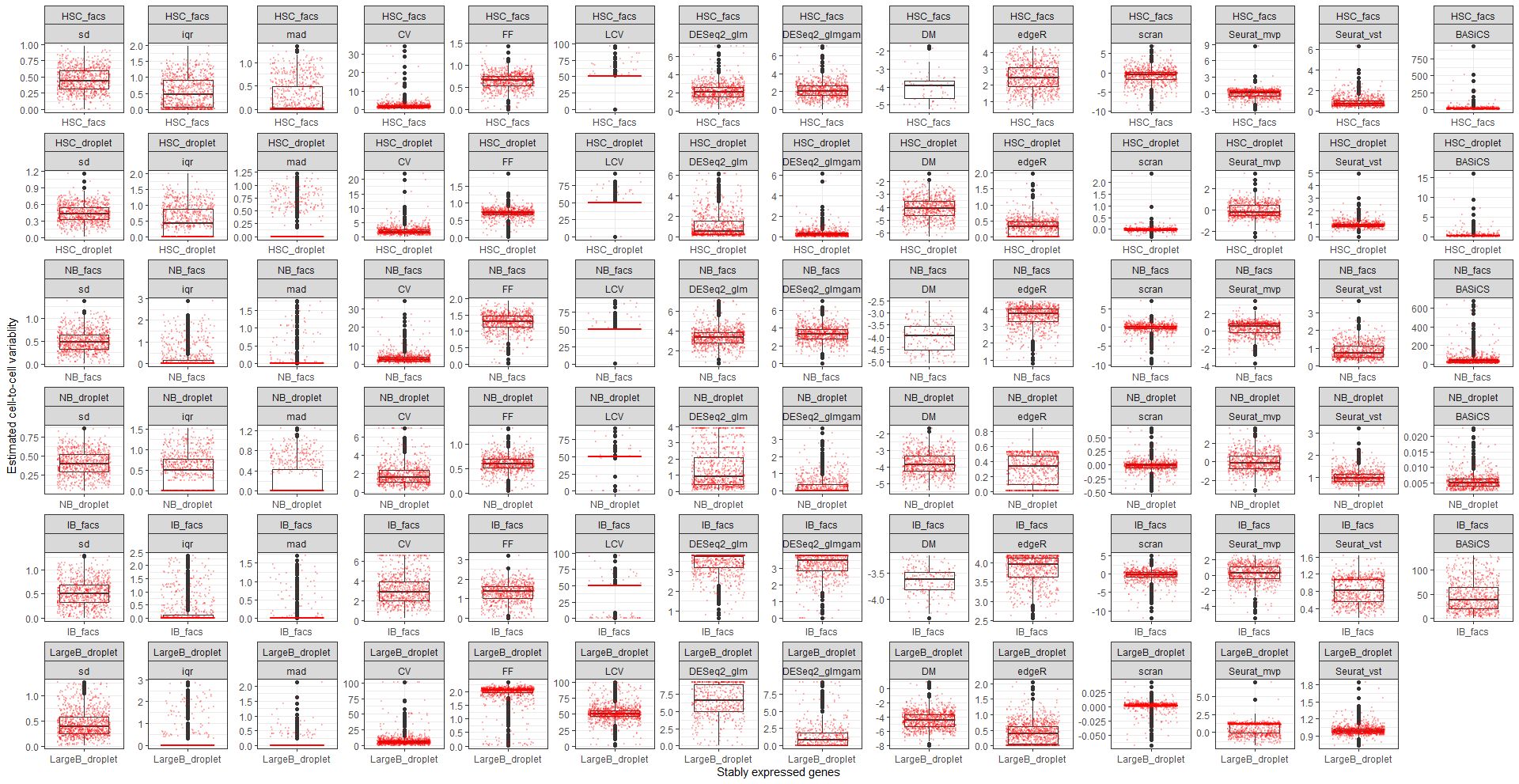
**E**

Supplementary figure 2: Plots for each metric between the estimated cell-to-cell variability and each aspect of data characteristics. Each dot in the plot represented a gene. Each row represented one dataset that had been tested. a) Dot plots for estimated cell-to-cell variability against the number of zero per gene across all cells. b) Dot plots for estimated cell-to-cell variability against the log mean expression. c) Dot plots for estimated cell-to-cell variability against available gene length. d) boxplots for each measured cell-to-cell variability with ribosomal genes coloured in red. e) boxplots for each measured cell-to-cell variability with stably expressed genes coloured in red.

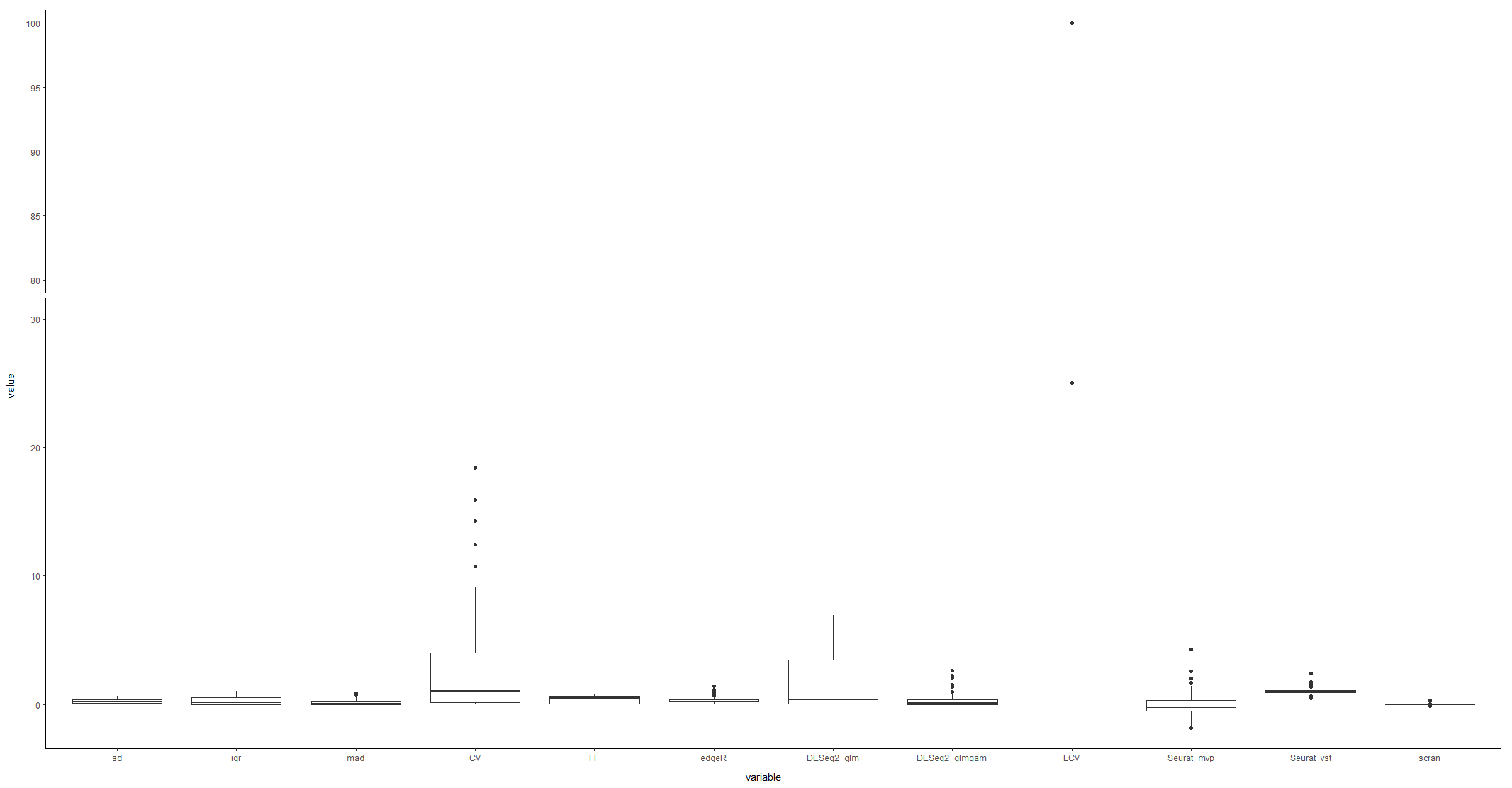

0.17

0.30

18.28

0.94

0.31

0.04

0.23

5.20

0.28

0.24

2.24

0.48

Supplementary figure 3: Metrics measured on the spike-in control dataset that only contained technical variability as shown in the boxplots. The measurement dispersion was calculated and listed for each metric in the figure. The lower dispersion value illustrated the capability of measuring biological variability rather than technical variability by a metric. DM and BASiCS were not included as required gene length and batch information, respectively.

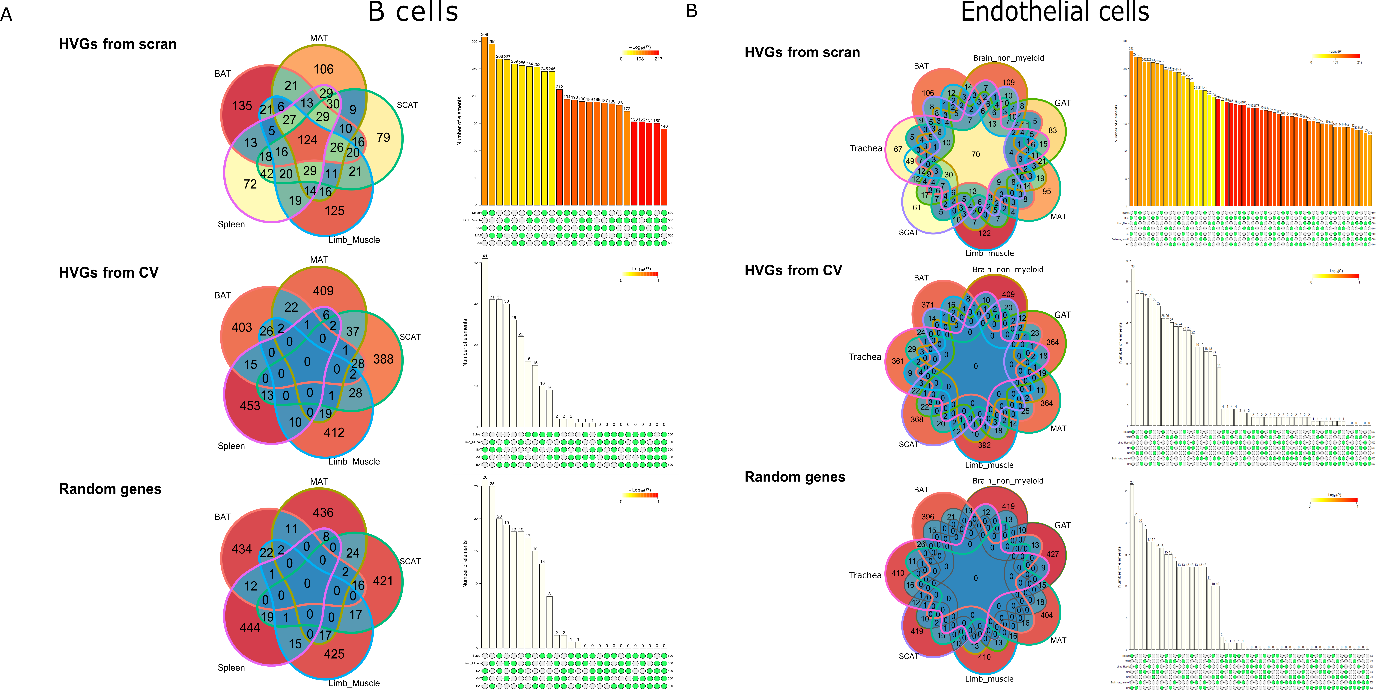

Supplementary figure 4: Venn diagrams showed the overlapped percentage of highly variable genes (HVGs) for B cell (A) and endothelial cells (B) measured by scran across multiple tissues (BAT: brown adipose tissue; MAT: mesenteric adipose tissue; SCAT: subcutaneous adipose tissue; GAT: gonadal adipose tissue;) and barplot showed the overlap of every listed with the colour indicated p-values. By further comparing with random sampling genes, the overlap of HVGs captured with random gene lists showed high similarities with CV but not scran.

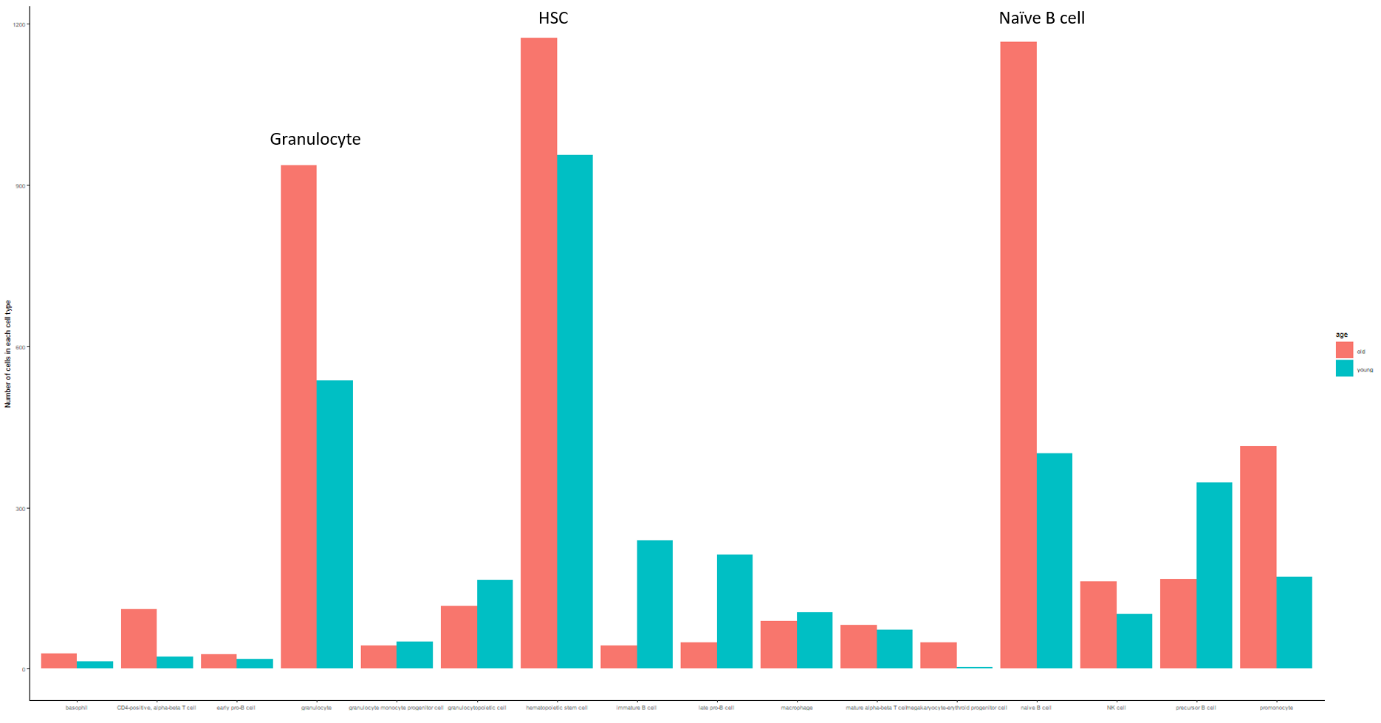

Supplementary figure 5: Barplot illustrated the number of cells in the marrow tissue in both young and old groups in TMS. To investigate the B cell lineages differentiation and aging, only related cell types with more than 100 cells in either age group were included, which are HSC, late progenitor-B, precursor B, immature B and naïve B cells.

Supplementary figure 6: Volinplot of gene expression variability for five cell types between young and old groups from Tabula Muris Sensis data. Out of five cell types, only precursor B
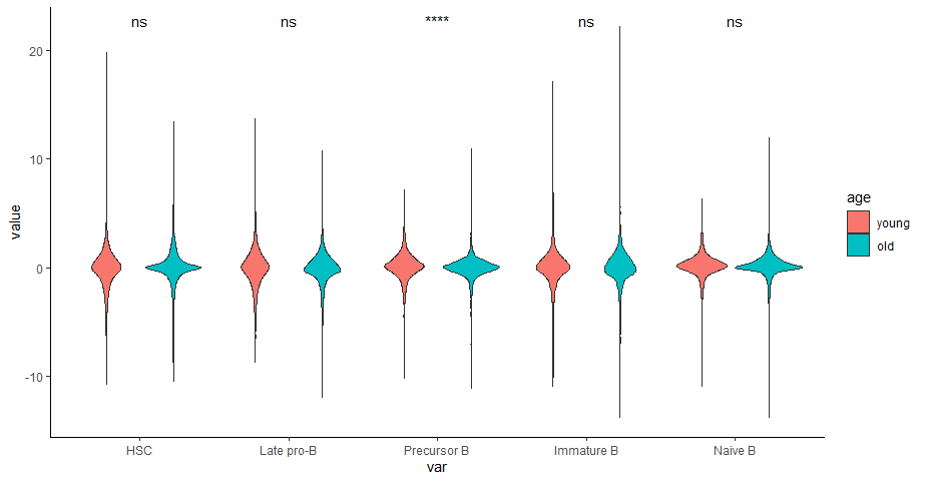
 cells showed a significant difference between young and old groups by performing the Wilcoxon ranked sum test. ns represented no significance and **** represented p.value < 0.0001.

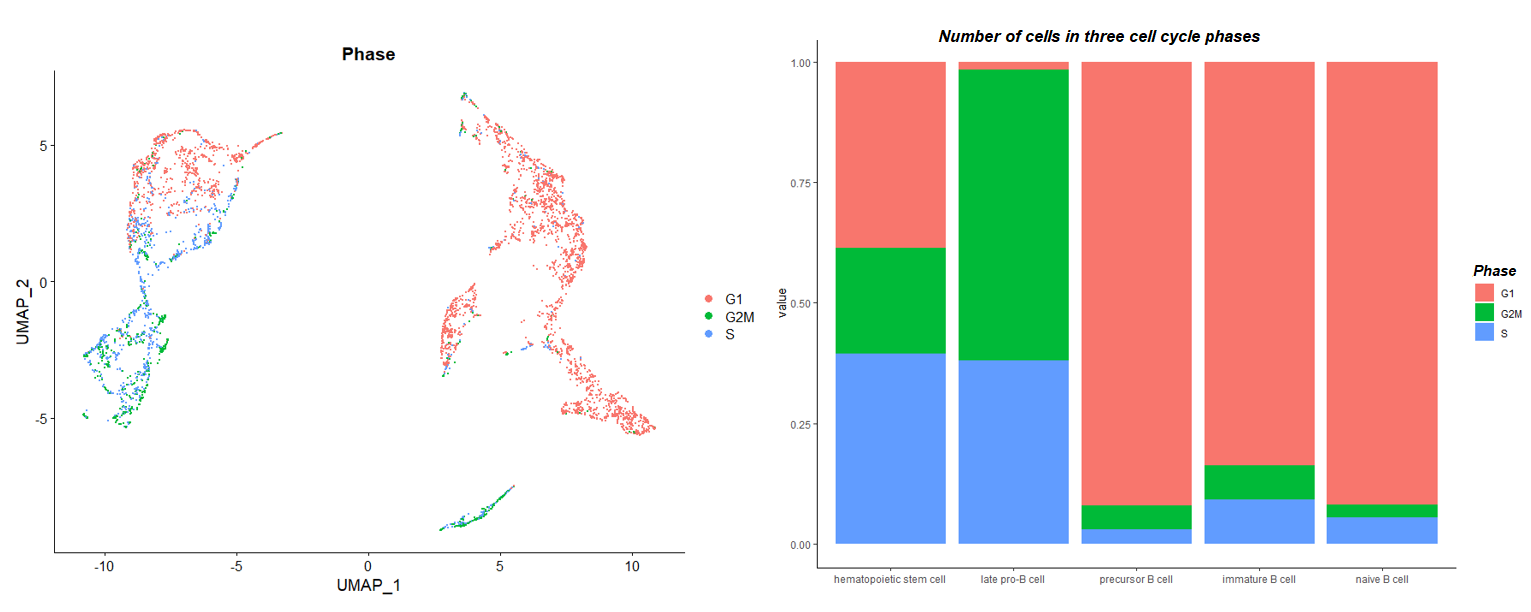

Supplementary figure 7: The cell cycle stages of HSC, late pro-B cells, precursor B cells, immature B cells and naïve B cells. Stages were assigned to each cell and coloured in the UMAP. Barplot showed the corresponding proportions of cells in the G1, G2M or S phase.

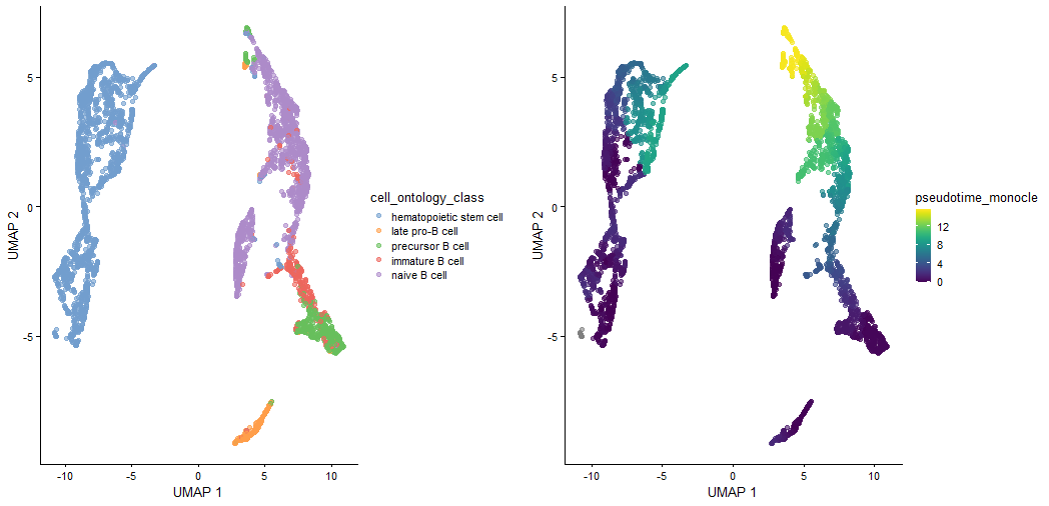

Supplementary figure 8: UMAP illustrated the pseudotime inference for each cell type after removing the age effect. The pseudotime trajectory matched the biological B cell differentiation, however, HSC and naïve B cells showed outspread pseudotime inference.

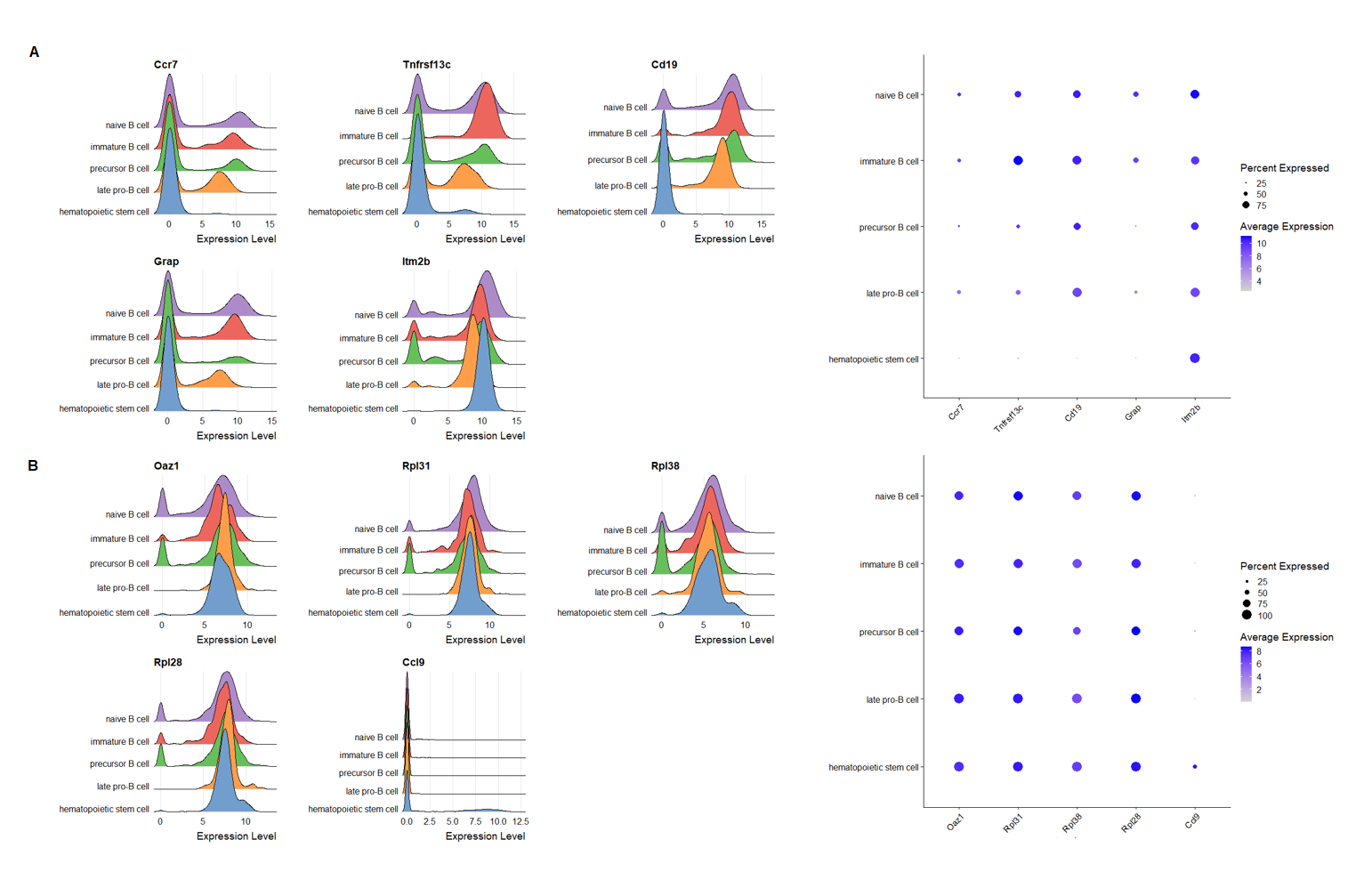
Supplementary figure 9: Details of the top 5 consistently variable genes and consistently stable genes along the B lymphocytes differentiation process. Ridgeplot of expression and dotplot of the percentage of expression in each cell type for A. consistently variable genes, B. consistently stable genes.

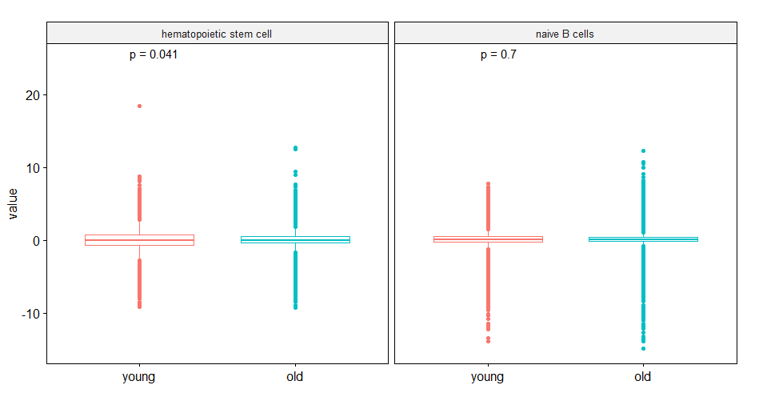
A

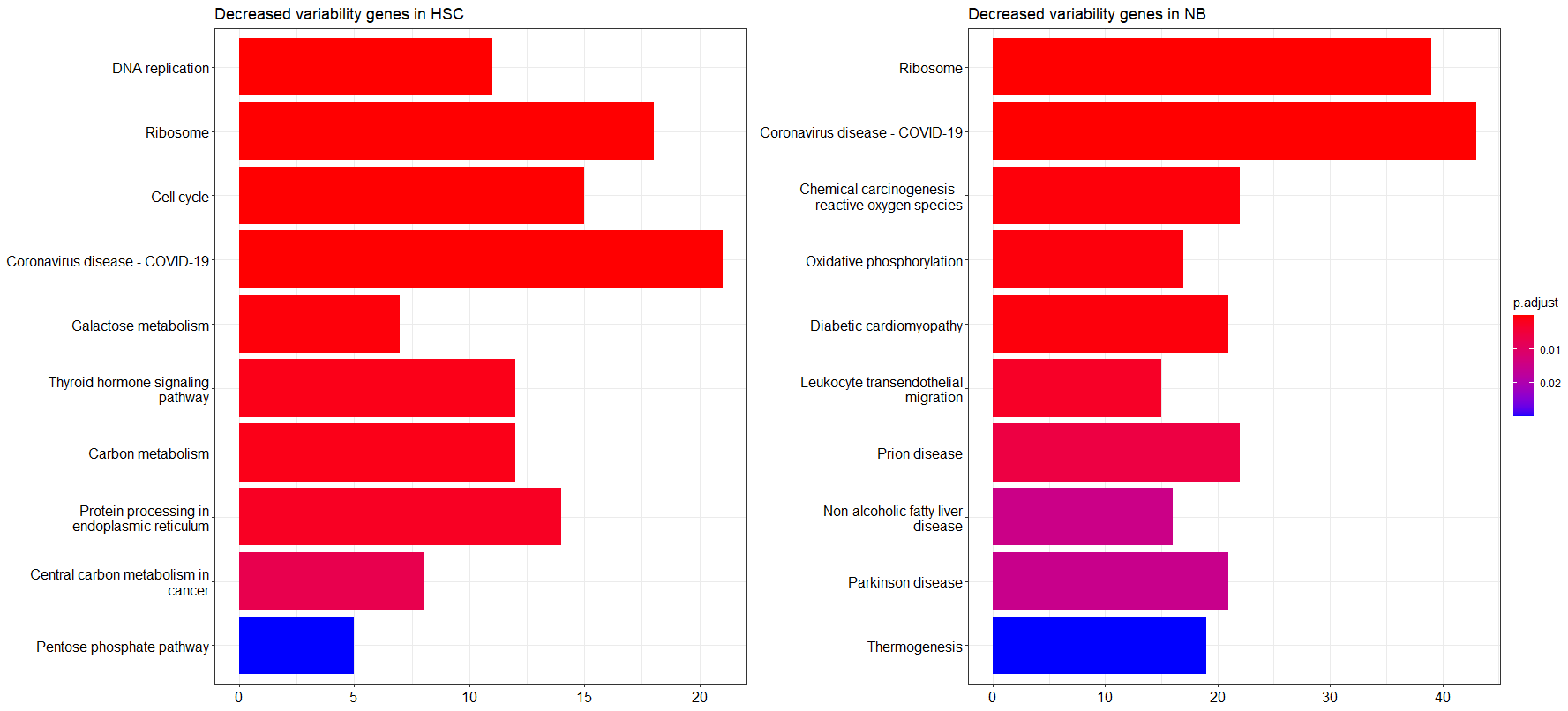
B	C

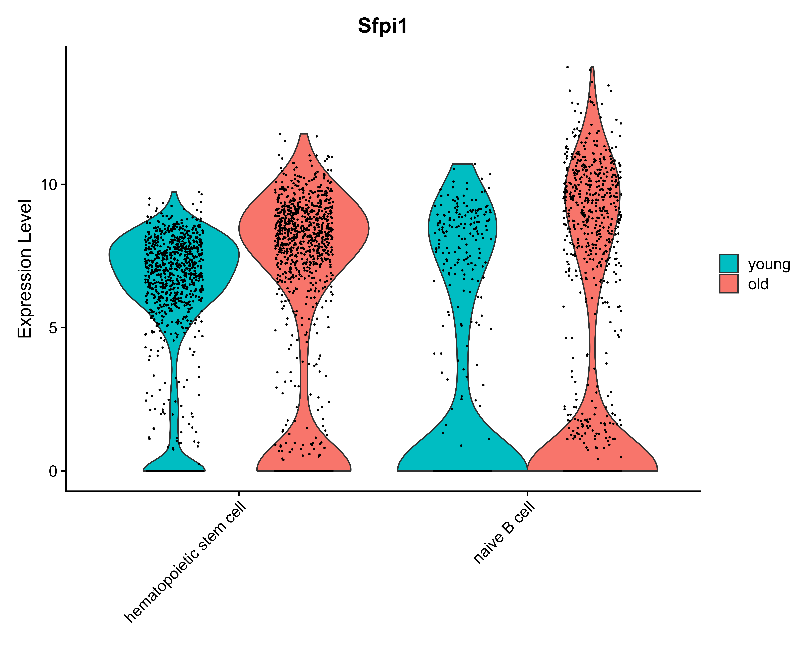

D

Supplementary figure 10: Cell-to-cell variability alterations in HSC and B lymphopoiesis in aging. A. overall distribution of measured variability in HSC and NB between young and old groups. p.value was calculated by the Wilcoxon test for each cell type. B. Top 10 significant pathways from KEGG databased based on significantly decreased variability genes from HSC, ordered by p.value. C. The top 10 significant pathways from KEGG databases based on significantly decreased variability genes from D. Violin plot highlighted the gene expression level of *Sfpi1* between young and old groups for HSC and naïve B cells.
